# Optimization of the South African indigenous fungal growth for the degradation of diclofenac sodium from water

**DOI:** 10.1101/329250

**Authors:** Teddy Kabeya Kasonga

**Affiliations:** Department of Environmental, Water and Earth Sciences, Tshwane University of Technology, Arcadia Campus, Private BagX680, Pretoria 0001, South Africa

**Keywords:** Degradation, fungal growth, diclofenac removal, NSAID, South African indigenous fungi

## Abstract

**Background and objectives:** The occurrence of endocrine-disrupting compounds (EDCs), pharmaceuticals and personal care products (PPCPs) or active pharmaceutical ingredients (APIs) and their risk assessment in the environment over a decade have become a real concern in various existing water resources. Microbial bioremediation of organic pollutants in wastewater is a key process in both natural and engineered systems. This study aimed to the use of green technology with South African indigenous fungi for the removal of diclofenac from water, which is an environmentally friendly process applied to manage water quality at large.

**Method:** The fungal growth was optimised in flasks, then the aerated and stationary batch flasks were run for 14 d and samples taken once daily in order to carry out the fungal removal efficiency of the most popular and anti-analgesic anti-inflammatory drug (NSAID) diclofenac sodium (DCF) from water. The five isolate South African indigenous fungal strains (ISAIFS) *T. longibrachiatum, T. polyzona, A. niger, M. circinelloides* and *R. microsporus* were then found to have a optimum growth in low nitrogen medium (LN-m) at temperature range of between 26.5°C to 31.5 °C and pH around 3 to 4.5.

**Results:** *Aspergillus niger* gave better growth and seemed thermotolerence than others. Glucose supply as well as physicochemical parameters such as pH and temperature have shown to have play a vital role on fungal growth in suspension liquid media. The best DCF degradation result obtained was 95% by *R. microsporus* in aerated batch flasks after 7 d followed by A. niger with 80% of DCF removal, while the only one white-rot fungi (WRF) of that isolate fungal group, *T. polyzona* did not give the best DCF elimination as expected for the same period.

**Conclusion:** Finally, the effectiveness of DCF elimination by each isolate South African indigenous fungal strain (ISAIFS) was found to be better than some traditional methods used in wastewater treatment plants, including: coagulation-flocculation, nitrifying and denitrifying and sewage treatment. These fungal species especially *R. microsporus, A. niger* and *M. circinelloides* can be used for the degradation of emerging pollutant in wastewater treatment plants.

## Introduction

Over the past decades, emerging micropollutants such as endocrine-disrupting chemicals (EDCs) including pharmaceuticals and personal care products (PPCPs) have become a matter of concern in both drinking water and wastewater treatment plants (WWTPs) (Rodarte-Morales *et al*., 2012a,b; Verlicchi *et al.*, 2012; Li *et al.*, 2014; Nam *et al*., 2015). The failure of WWTPs to completely remove a wide range of PPCPs from wastewater results in the contamination of the aquatic environment as their concentrations are able to increase from few ng/L to 15μg/L, levels which are enough to generate acute and chronic toxicity (Jones *et al.*, 2005; Li *et al*., 2014; Verlicchi *et al.*, 2015), with risk of degrading of environmental aesthetics as well as compromising the ecosystem services (Whitehead and Crossman, 2012).

Among those EDCs, PPCPs and APIs or PhACs, which constitute a real environmental threat, there is an analgesic painkiller and most popular non-steroidal anti-inflammatory drug (NSAID). Diclofenac sodium or potassium (DCF), 2-[(2, 6-dichlorophenyl) amino] phenyl acetic acid monosodium salt are the most frequently identified pharmaceuticals in different aquatic environments (Zhang *et al*., 2017). DCF was first introduced in Europe in 1973 (Daniels *et al.*, 2016), and has been available in several different topical formulations, such as DCF diethylamine gel 1.16%, DCF 1% gel, DCF DMSO lotion, MIKA DCF spray 4% gel and DCF epolamine (DCF hydroxyethyl-pyrrolidine) patch (Hui et al., 1998; Abraham *et al.*, 2014; Moreira and Liu, 2017). DCF has been also available at diverse names including Novapirina, Orthofen, Feloran (Gryshchenko, 2017), Dichlofenal, Voltaren (Moreira and Liu, 2017), Anthraxiton (Vieno and Sillanpää, 2014), Delphimix (Stöhr *et al,.* 1980), Allvoran, Arthrotec (Agrawal *et al.*, (1999), Vonafec, Vurdon (Goci *et al.*, 2014), GP45840 (Lepore *et al.*, 2017) and Cataflam (Shaw *et al.*, 2000).

DCF has analgesic, anti-inflammatory and antipyretic properties. As most of NSAIDs, it inhibits selectively the cyclooxygenase isoenzyme 1 and 2, and acts upon other inflammatory pathways (Strom *et al.*, 1996), by decreasing the production of the prostaglandins at a site of disease or injury wich are responsible for inflammatory and pain, (Ikehata *et al*., 2006). Therefore, NSAIDs have peripheral effects and operate in the spinal cord and central nervous system (Gan *et al.*, 2010; Hoy, 2016). Unfortunately, NSAIDs use to induce acute kidney injury as they can be depleted in volume and gastrointestinal bleeding (FitzGeralg, 2004), to generate hepatitis (Gryshchenko, 2017) and might increase the risk of cardiovascular event as well as liver damage (APS, 2008). Thereby the removal of DCF from water has been a focus of various studies. The occurrence of DCF has been reported by Zhang *et al*. (2008) and Rodarte-Morales *et al.*, (2011), in diverse water bodies such as WWTP effluents, groundwater, surface and drinking water. In South Africa, the estrogenic activity of these EDCs has been detected in Cape Town and Pretoria drinking water (Van Zijl *et al.*, 2017). Anderson *et al*. (2005) have mentioned the responsibility of DCF sodium for the catastrophic decline of three species of vulture populations in South Africa. DCF has been detected at the concentration of 9.69μg/L in Kwazulu-Natal Province river water (Madikizela and Chimuka, 2017).

Additionally, biological and physicochemical treatments such as coagulation-flocculation (Carballa *et al.*, 2005), sewage treatment (Kosjek *et al.*, 2007), nitrifying and denitrifying bioreactor (Suárez *et al.*, 2005), membrane bioreactors (Reif *et al.*, 2008), have been used for the removal of NSAIDs particularly DCF in environment. It has also been reported by Cherik *et al.* (2015) that in the biological process such as biodegradation by activated sludge and membrane bioreactor, the presence of DCF can affect the bacterial community in sludge and influence the efficiency of the biodegradation process. Therefore, DCF is considered as one of the recalcitrant pharmaceuticals barely removed in WWTPs with respect to their chemical structures (Banaschik *et al.,* 2018). Furthermore, as one of the most abundant NSAIDs found in wastewater and considering its physicochemical parameters such as molecular weight (MW=296.2 g/mol), solubility in water (S_w_=2.4 mg/L), vapor pressure (P_v_=8.10-6 Pa), dissociation constant (pKa=4.0-4.5) and octanol-water partition coefficient K_ow_ (logK_ow_= 4.5-4.8) (Ziylan and Ince, 2011; Suárez *et al.*, 2008; Scheytt *et al*., 2005; Lai *et al*., 2000), the simple removal of DCF from environment might be unexpected using conventional wastewater treatment technology. Because of the pKa values, the ineffective adsorption process for acid drugs including DCF has been reported by Nikolaou *et al.* (2007). However, the DCF low solubility in water (S_w_) which expresses its hydrophobicity coupled to its low vapor pressure (P_V_) contribute to decrease the DCF aqueous phase concentrations as well as its negligible volatilization from water. Even though in view of its relative high logK_ow_ values, the sorption on sediment or soil might be an important factor of DCF removal. Madikizela and Chimuka (2017) have reported from Kwazulu-Natal Province river water that, due to their polar nature, pharmaceuticals such as DCF, ibuprofen and naproxen do escape the wastewater treatment process and consequently contaminate water. Therefore, the bioremediation may be considered remain as the DCF expected potential elimination pathway mechanism from wastewater.

Furthermore, since fungi have been found to possess an uncommon bioremediation ability of various hydrocarbons (Rodarte-Morales *et al.*, 2011; Golan-Rozen *et al.*, 2015; Barnes *et al.,* 2018). Because of their modifying enzymes (Majeau *et al.*, 2010), their utilization in biodegrading a number of the recalcitrant compounds might reduce some stress of the adjacent water bodies (Majeau *et al.*, 2010; da Silva *et al.*, 2017). Therefore, the use of other fungal species such as ascomycetes and zygomycetes such as t the isolated South African indigenous fungal strains could be a valuable alternative in the removal of NSAIDs from wastewater. Hence wastewater pollutants in general can be adsorbed and degraded by dead and living fungal biomass as adsorbent support. This doesn’t affect the microbial physiology as reported by Takey *et al.* (2014).

The present study aimed to stablish the optimum growth of the South African indigenous fungal growth in suspension liquid media and investigate their efficiency in removing analgesic and non-steroidal anti-inflammatory drug (NSAID) diclofenac sodium from synthetic wastewater. The following five (5) South African indigenous fungal strains (ISAIFS) isolated from our previous: were used to achieve the aim of this study: *A. niger, M. circinelloides, T. longibrachiatum, T. polyzona* and *R. microspores.*

## Materials and methods

### Preparation of the fungal culture media

The chemicals and reagents used in the present study series were of analytical grade purchased from Sigma Aldrich, South Africa and Minema Chemicals, South Africa

#### The solid medium D for fungal culture and maintenance

The medium D contained 40g of D-(+)-glucose anhydrous, 15g of agar and 10g of peptone protease per liter. After sterilisation by autoclave, the medium was transferred into sterile Petri dishes and maintained at ambient temperature in aseptic conditions prior to use. The five South African indigenous fungal strains (ISAIFS) - *T. longibrachiatum, T. polyzona, A. niger, M. circinelloides* and *R. microspores* were cultured in the solid medium D for 7 d at 30±1.5 °C. Thereafter, the fungal strains were maintained in medium D stored at 4°C.Preparation of liquid media for fungal mycelial growth

#### Preparation of the liquid media for the fungal mycelial growth

##### Preparation of the low nitrogen medium (LN-m)

The liquid medium LN-m was prepared as described by Tien and Kirk (1988) for inoculums production. Briefly, it comprised 100mL of Basal III media, 100mL of 0.1M of *trans* aconitic acid (pH 4.3) (Sigma Aldrich, SA), 10g of D-(+)-glucose anhydrous (Sigma Aldrich, SA), 0.2g of Ammonium tartrate dibasic (AT) (Sigma Aldrich, SA) and 60 mL of trace elements. The solution was then autoclave at 120°C for 30 min. Finally, 1mL of sterile thiamine chloride solution (1mg/mL), was added to the cooled media. It is important to note that 100 mL of the sterile 4mM 3,4-dimethoxybenzyl alcohol (96%, MINEMA Chemicals, SA) (Veratryl alcohol: VA) was added on the third day of incubation of the microorganisms to stimulate the enzyme production and to moderate the enzyme inactivation in the media (Koduri and Tien, 1994; Zhang and Geißen, 2010). *Trans* aconitic acid was used as buffer instead of 2,2-dimethylsuccinate provided in Tien and Kirk (1988) medium recipe. The final pH of the LN-m (±4.3) was reached by using NaOH or HCl 0.1M. The measurement of the pH value of the media was ascertained using a HACH multi HQ-40d pH-meter provided with HACH-CDC401 electrode.

##### Preparation of the Puig (2012) liquid medium (PL-m)

The liquid medium (PL-m) was prepared according to Piug (2012) and contained: 10g of D-(+)-glucose anhydrous, 5g of peptone protease, 2g of KH_2_PO_4_, 0.5g of MgSO_4_.7H_2_O, 0.1g of CaCl_2_, 10mL of trance element solution and 2mg of the sterile thiamine chloride (1mg/L) per liter.

##### Preparation of the Basal III solution

The *Basal III medium* containing 20g of KH_2_PO_4_, 5g of MgSO_4_.7H_2_O, 1g of CaCl_2_ (MINEMA Chemicals, SA) and 100 mL of trace elements solution per litre was prepared and stored in the refrigerator as described by previous investigators (Kirk *et al*., 1986; Tien and Kirk, 1988).

##### Preparation the trace elements solution

The reagents used for the preparation of the trace elements solution were of analytical grade and purchased from MINEMA Chemicals South Africa. In brief this solution contained 1.5g of Nitrilotriacetic acid pH 6.5 (adjust pH using NaOH or HCl, 1M), 3g of MgSO_4_.7H_2_O, 0.5g of MnSO_4_.H_2_O, 1g of NaCl, 0.1g of FeSO_4_.7H_2_O, 0.1g of CoCl_2_.6H_2_O, 0.1g of ZnSO_4_.7H_2_O, 0.1g of CuSO_4_.5H_2_O, 10mg of Al_2_(SO_4_)_3_.18H_2_O, 10mg of H_3_BO_3_ and 10mg of Na_2_MoO_4_.2H_2_O per litre. It was prepared and then stored in the refrigerator as indicated by previous investigators (Kirk *et al.*, 1986; Tien and Kirk, 1988).

### Preparation of fungal mycelium

The biomass of fungal mycelium was determined using a spectrophotometer (HACH model DR 6000TM, Colorado, USA) which was equipped with 10 mm glass cells for the measurement of absorbance. The volume of 10 mL of fungal spore solution at 0.5cm^−1^ of absorbance at 650 nm corresponding to 2.5×10^6^/mL fungal spores was inoculated in 90 mL of LN-m or PL-m media. The fungal mycelia obtained after 5 d of spore incubation at 30±1.5°C were mixed using the sterilized blender. The resultant mycelium solution was used to spike single fungal batch flask for the pollutant degradation study as described by Tien and Kirk (1988).

### Batch flask medium composition

Each 500 mL flask used as laboratory batch reactors containing 200 mL of synthetic DCF contaminated water was prepared as followed: 10% (v/v) of fungal mycelial inoculum and 0.2g of ammonium tartrate dibasic (AT) or 0.2g of ammonium sulfate (Sigma Aldrich, South Africa). The sterile Tween^®^ 80 (polysorbate 80) (1mL) was added to each aerated batch flask as surfactant for fungal pellet formation as reported by Liu and Wu (2012). Tween^®^ 80 was also used for fungal extracellular enzymes protection from mechanical degradation (Tien and Kirk, 1988). The control batch flasks were performed without fungal mycelium.

### Evaluation and optimisation of fungal growth in liquid media

#### Fungal growth evaluation

The assessment of the fungal growth in suspension liquid media was performed using filtration method and by weighting the fungal dry mass from batch flask (Patel *et al.*, 2017; Yamamoto *et al.*, 2017). Briefly, prior to proceed to the filtration of the fungal suspension, the labeled Whatman (0.4 to 0.5 μm) filter papers was weighed. Thereafter, the filter paper containing the fungal biomass was covered with the aluminum foil and subjected to dry into the oven for 48 h at 40°C. From the oven, the filter paper was allowed to cool in the desiccator and the fungal biomass was calculated based on the difference values obtained from the weight of the filter paper before and after filtration of the fungal suspension. Flasks [100mL of LN-m or PL-m and 200mL of batch flasks] were taken to weigh fungal mass after 3, 6, 9, 12 and 15 d of incubation. Experiments were run six (6) times for the first batch flasks and in triplicate for the last batch flasks

#### Effect of nutrient on fungal growth in selected liquid media

The fungal growth study performed in two liquid media [LN-m and PL-m] at 30±1.5°C for 15 d. Fungal mycelium flasks were taken after 3, 5, 9, 12 and 15 d and the dry mass was obtained as described above.

#### Effect of temperature on fungal growth in LN-m

The influence of temperature on the fungal growth in the LN-m was evaluated at 28±1°C, 30±1.5°C and 37±1°C at a starting pH 4.3 for 15 d and the fungal dry mass was evaluated after each 3 d. The fungal thermotolerance study in LN-m was performed at nine (9) different temperatures (20±1, 25, 30±1.5, 35, 40, 45±1, 50, 55±1.5 and 60°C) and the fungal dry mass was obtained after 5 d of incubation.

#### Effect of pH on the fungal growth in LN-m

The effect of the starting pH on fungal growth in LN-m was carried out at 30±1.5°C for 5 d of incubation as described by Tien and Kirk (1988). Different values of pH were considered (2.3, 3.1, 4.3, 5.5 and 6.2) at the onset of fungal spores’ inoculation. The final pH values were also recorded after 5 d of incubation and the fungal dry mass was calculated as described above.

#### Effect of air supply on fungal growth in batch flask

The fungal growth assessment in batch flasks containing 200 mL of media were conducted at room temperature in both stationary batch flask (StBF) and aerated batch flask (ABF). The aerated batch flasks were supplied with humidified air through water flask. The fungal dry mass was calculated as described above.

#### Effects of nutrient on fungal growth in batch flasks

Different nutrients [glucose 0g/L versus 10g/L, ammonium tartrate dibasic (AT) and ammonium sulfate, 0g/L against 0.2g/L] were added in batch reactors at room temperature in order to carry out the effect in fungal growth.

#### Effect of sterile condition and daylight on fungal growth in StBF

The non-sterile StBF was conducted with tape water in opened 500 mL bottles, while sterile batch was performed in closed sterile 500 mL bottles using DDW as described in Section 2.3. The StBF totally covered with aluminum foil were used to perform the batch without light against the uncovered batches.

### DCF analytical method and optimisation of its fungal removal efficiency

#### DCF analytical method

At room temperature, the monitoring of the standard diclofenac sodium (DCF 99.8%, Sigma Aldrich, South Africa) spiked at the concentration of 10 mg/L in batch flasks was carried out spectrophotometrically based on the method described by Matin *et al.* (2005) with slight modification. Briefly, in this experimental study, the nitric acid was at 55% instead of 63% as described by the authors. The reaction between DCF and concentrated nitric acid 55% (2:1, v/v) resulted in a yellowish complex solution at an absorbance of 380 nm. A blank was also included using a mixture of water and nitric acid 55%. Samples were taken daily during the study period of 14 d to assess the DCF removal efficiency at room temperature (27±1°C). Experiments were run in triplicate. The effects of air supply, nutrients, non-sterile conditions and daylight on DCF removal by fungi were conducted as described above.

#### Effect of fungal biomass concentration on DCF removal efficiency

To assess the effect of the fungal biomass on DCF removal efficiency, fungal mycelium solution was inoculated in the StBF at different concentrations (5, 10, 20, 30, 40 and 50% v/v). The experiments were run for 5 d in individual fungal strain batches to evaluate the DCF removal efficiency in term of fungal biomass increases.

### Statistical analysis

The data analysis in the present study was done using vegan package in R (ver. 3.4.1) statistic software to perform the data analysis (Oksanen *et al.*, 2013) for comparing fungal biomass growth of identified indigenous fungal strains in liquid media as well as the DCF removal efficiency from different performed batch flasks.

## Results

### Fungal growth in liquid media

Fungal mycelium growth in suspension media

#### Effect of selected media

Figures 1 and 2 illustrate the growth of selected South African indigenous fungi in liquid media at 30±1.5°C. All data are represented in average values of 6 experimental study series.

**Figure 1:**
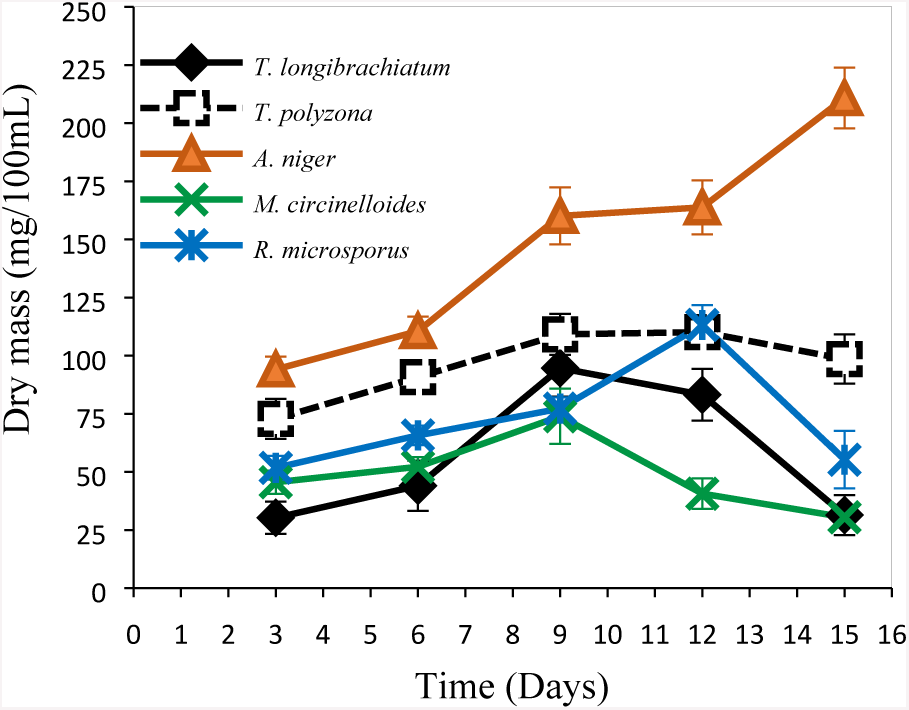
Fungal growth study at 30±1.5°C in LN-m with initial pH 4.3.

**Figure 2:**
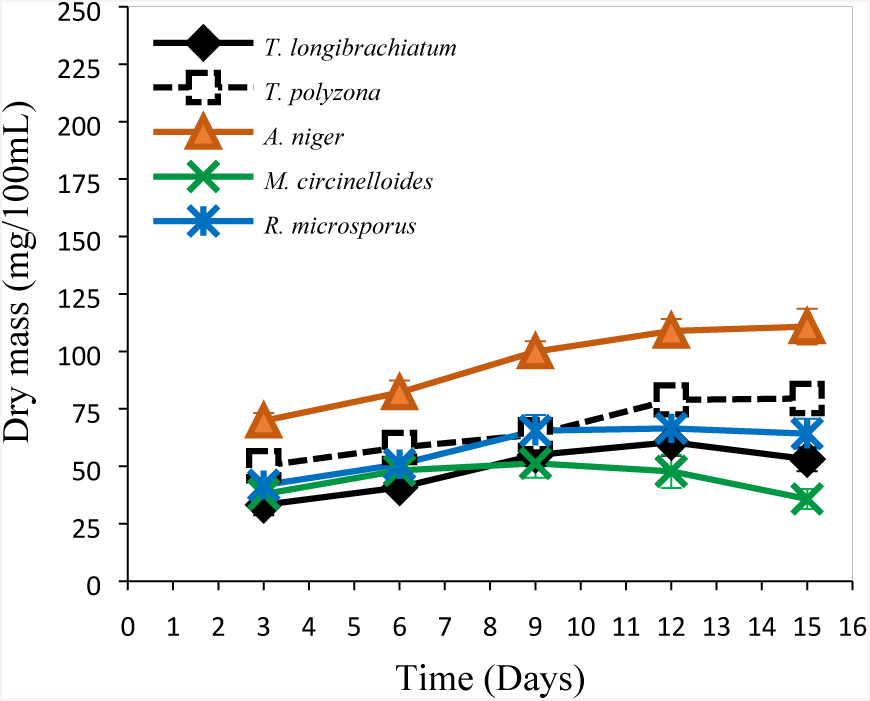
Fungal growth study at 30±1.5°C in PL-m with initial pH 4.3.

The LN-m seemed to give, in general, a good fungal growth compare to the PL-m. *A. niger*‘s biomass is higher (*p* < 0.001) than others fungi regardless of the media as shown in Figures 1 and 2. *T. polyzona* was the second to growth better followed by *R. microsporus* while *T. longibrachiatum* and *M. circinelloides* were the last. In addition, regardless of the media, *A. niger* was still growing until the 15 d, whereas other fungal strains were dying/decreasing from the day 9 or 12. The good growth in LN-m compare to PL- m (*p* < 0.001) might be due to the media compositions, except the contribution of micronutrients added as trace elements in the media. According to the media composition, the C/N ratio is 50:1 with ammonium tartrate dibasic (AT) as nitrogen source in LN-m and 7% of trace elements as well as *trans* aconitic acid and veratryl alcohol (VA), while the C/N is about 2:1 in PL- m using peptone as nitrogen source (Jonathan and Fasidi, 2001) and 1% of trace elements without VA neither *trans* aconitic acid. The higher C/N ratio might favour the fungal growth in LN-m. A higher C/N ratio gave a maximum fungal biomass as the case in LN-m.

A significant media effect (*p* < 0.001) was also observed among fungal strains compare to *A. niger*. Thus, at the third and sixth days, the fungal biomass (mass in mg/100mL) were respectively 30.26±6.8 and 44.02±10.5 for *T. longibrachiatum*, 72.8±8.5 and 90.8±4.9 for *T. polyzona*, 94.1±5.34 and 110.55±6.37 for *A. niger*, 45.55±5 and 52.2±3.35 for *M. circinelloides* then 51.75±5 and 65.7±4.25 for *R. microsporus* in LN-m while low values were collected in PL- m, meaning 33.22±4.55 and 40.8±3.25 for *T. longibrachiatum*, 50.21±3.4 and 58.2±4.3 for *T. polyzona*, 69.8±3.45 and 82.1±5.2 for *A. niger*, 37.9±4.5 and 48.2±3.55 for *M. circinelloides* and then 41.8±3.8 and 50.9±4.25 for *R. microsporus*.

In addition, considering the fact that the macro elements concentrations were equals (2 g/L of KH_2_PO_4_, 0.5 g/L of MgSO_4_.7H_2_O and 0.1 g/L of CaCl_2_), the fungal growth could have been promoted by trace elements contribution. The LN-m contained almost 7% of trace elements solution against 1% in PL-m. These elements as well as the addition of exogenous VA are enhancing extracellular fungal enzyme activities and different metabolisms. Therefore, *A. niger* gave better growth in LN-m (*p* < 0.001) followed in order by *T. polyzona* and *R. microsporus,* then came *T. longibrachiatum* and *M. circinelloides.*

#### Effect of temperature in LN-m

The temperature affects growth rate, nutritional requirements, regulation mechanisms of enzymatic reactions, metabolism and cell permeability in a microorganism, its influence in fungal suspension growth was carried out in LN-m at 28±1°C and at 37±1°C as given in the following Figures 3 and 4, comparing to 30±1.5°C shown in Figure 1 above.

**Figure 3:**
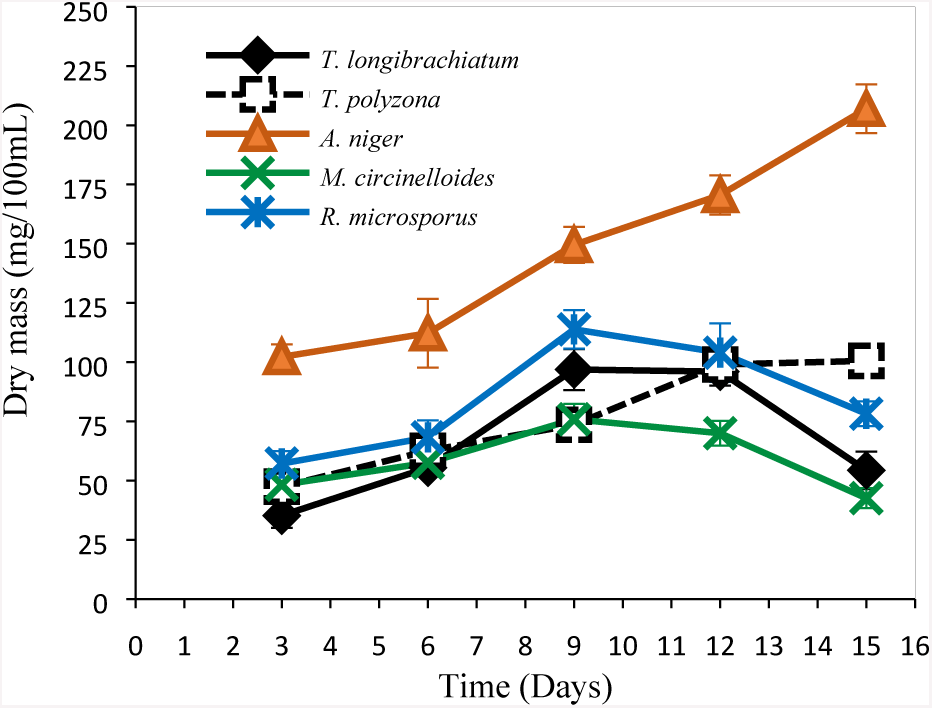
Fungal growth study at 28±1°C (in LN-m, initial pH 4.3)

**Figure 4:**
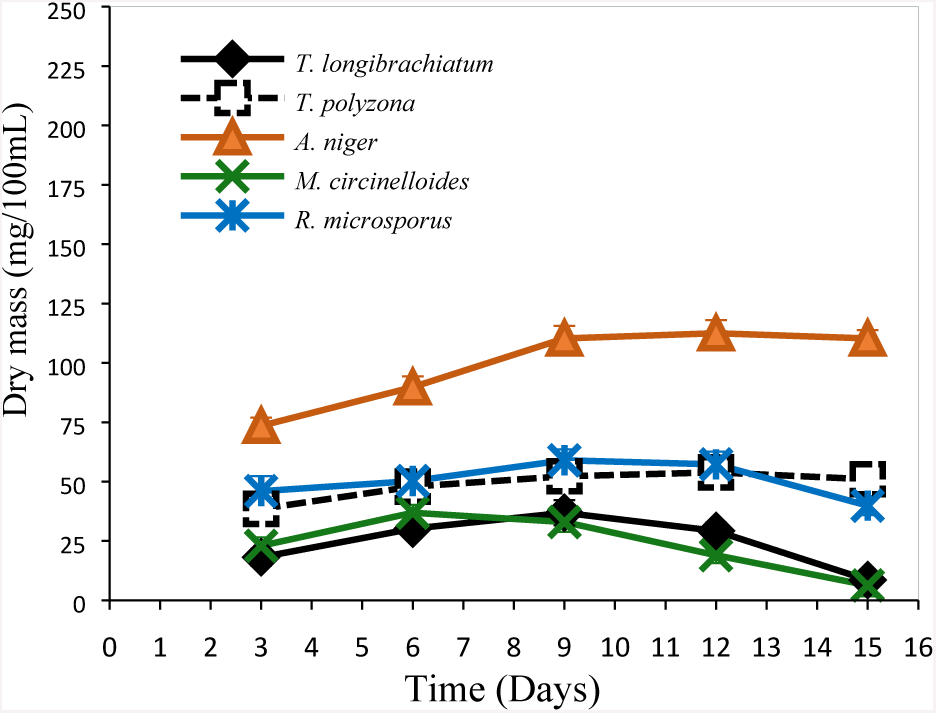
Fungal growth study at 37±1°C (in LN-m, initial pH 4.3)

Considering the three temperature ranges as shown in Figures 1 (30±1.5°C), Figure 3 (28±1°C) and in Figure 4 (37±1°C), the fungal biomass seemed to be affected by the increasing of the temperature from day 9, with *M*. *circinelloides* as the most affected from day 6 at 37±1°C. The best fungal growth has been observed in the range of the temperature between 28±1°C°C to 30±1.5°C. Thus, the fungal destruction in relation to time appears important with the increasing of the temperature. However, *A. niger* was more resistant and gave the best growth regardless of the temperature. Therefore 30±1.5°C could be considered as working temperature for mycelium growth means inoculum preparation and proliferation within 5 d as recommended by Tien and Kirk (1988). Regardless of working conditions (including pH, culture media or temperature), fungi keep on growing after 5 d. However, at 37±1°C, *M. circinelloides* biomass started decreasing from day 6, while *A. niger* seemed to be affected from day 12.

#### Fungal thermotolerance study in LN-m

Figure 5 gave the fungal thermotolerance recorded at different temperatures. The temperature as a physicochemichal parameter effected fungal growth as well as their enzymatic dynamic. The ISAIFS‘s growth was temperature dependent in liquid media (Figure 5). Regardless of the fungal species, the increase of the temperature affected the fungal growth. The suitable temperature fungal growth was found to be in the temperature range of 25°C to 35°C with almost an optimum at 30±1.5°C, except *R. microsporus* which gave the optimum at 25°C. However, *A. niger* was found to be more thermo resistant compare to other isolate fungi (*p* < 0.05). At 45±1°C for example, its dry fungal biomass collected was 77.1±5.42 mg/100mL, against 19.2±2.55 mg/100mL for *T. longibrachiatum*, 24.3±5.01 mg/mL for *T. polyzona*, 20.2±2.99 mg/mL for *M. circinelloides* and 31.1±5.12 mg/mL for *R. microsporus*. Even though the growth of all the isolated fungal strains was found to be affected from 30°C, *T. polyzona* appeared to be more affected. A decrease of 39.5% and 82.1% were recorded in its fungal biomass, respectively when the temperature increased from 30°C to 35°C and from 30°C to 50°C.

**Figure 5:**
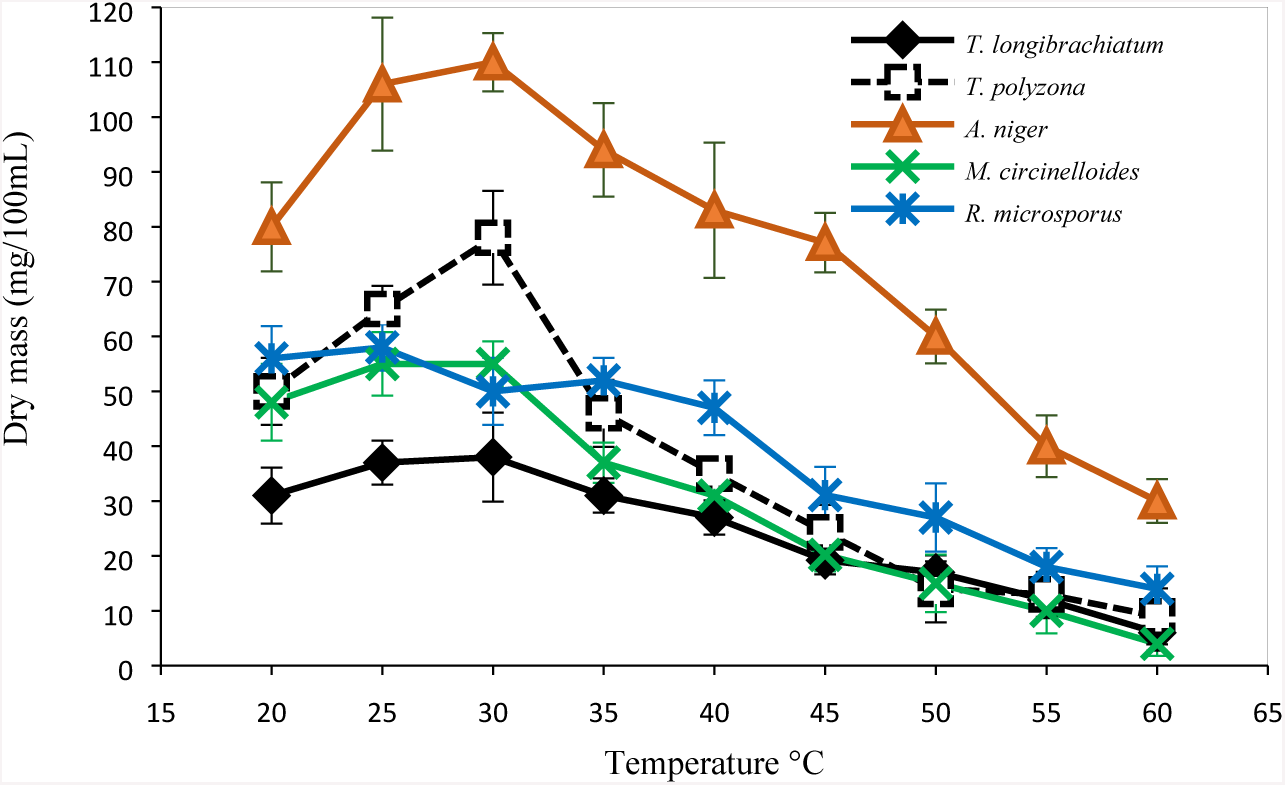
Temperature effect on fungal growth in LN-m.

#### Effect of starting pH of LN-m

In addition, Figure 6 below gives the pH effect of fungal growth in 100mL of inoculum after 5 d of incubation in LN-m at 30±1.5°C. This time is considered to allow fungal spores to growth for mycelium formation to allow for further inoculation in batch bioreactors. This experiment was performed in triplicate for each fungal strain.

**Figure 6:**
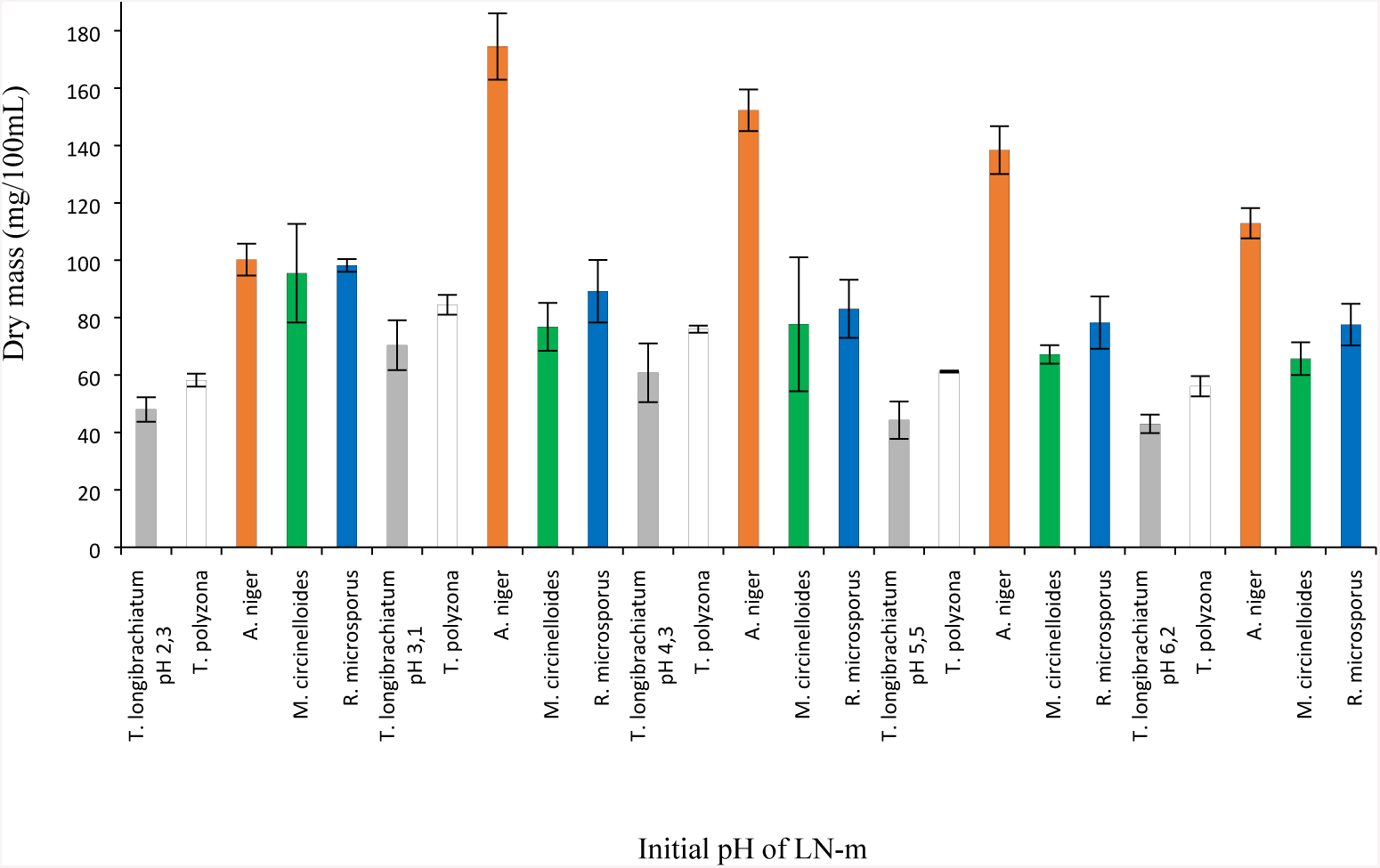
Effect of initial pH on fungal growth in LN-m at 30±1.5°C.

From the different selected values of initial pH: 2.3; 3.1; 4.3; 5.5 and 6.2, the fungal biomass achieved after 5 d was lower in the flasks with high starting pH, except for *M. circinelloides* (95.5 mg/100mL) and for *R. microsporus* (98.2 mg/100mL). The higher fungal growth was found to bet at the starting pH 3.1 for *A. niger* (174.5±11.55 mg/100mL), *T. polyzona* (84.5±3.45 mg/100mL) and *T. longibrachiatum* (70.4±8.71 mg/100mL). From the pH 3, the *A. niger* growth was significantly high compare to others regardless of pH (*p* < 0.001). These biomass values decreased with the increase of the pH. However, the fungal biomass obtained at pH 4.3 (152.3±7.25 mg/100mL for *A. niger*, 76.0±3.45 mg/100mL for *T. polyzona* and 60.8±10.22 mg/100mL for *T. longibrachiatum*) are not bad at all, at the time the values still at the same average with those at pH 3.1. Besides, the pH 4.3 was recommended by Tien and Kirk (1988) for enzyme production. Thereby, after 5 d, fungal mycelial hyphae were still growing in suspension media. This means that, at the collecting day of fungal mycelium to inoculate the batch cultures, fungal strains were still growing and producing metabolites.

#### pH evaluation in LN-m

The physico-chemical parameters including pH and temperature were monitored while fungal mycelium were growing in LN-m after incubation of 5 and for 15 d as showed respectively in Figure 7 and Figure 8 below. The Figure 7 gave the final pH after 5 d of incubation from different starting pH indicated in first column.

**Figure 7:**
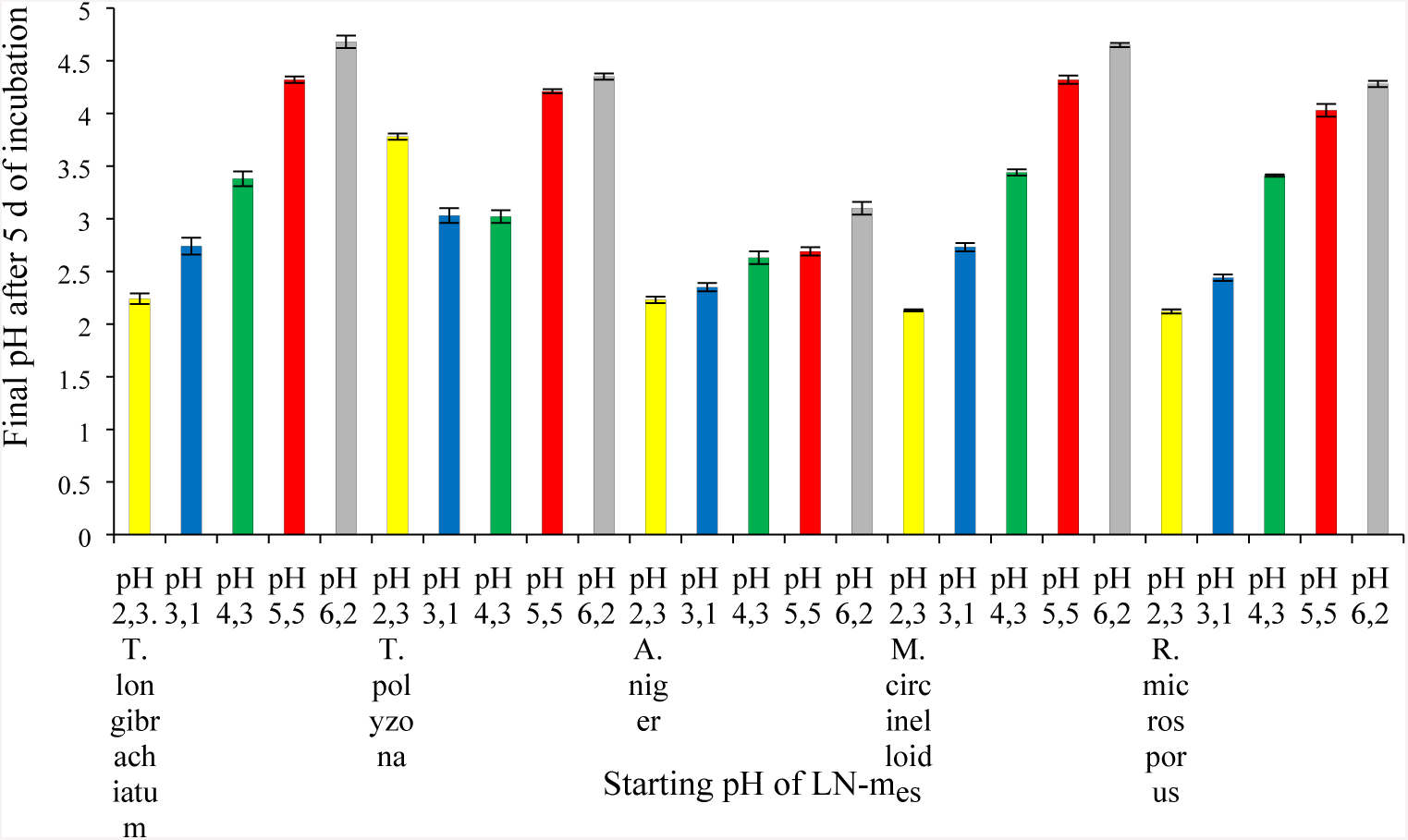
Final pH values of fungal LN-m after 5 d of incubation at 30±1.5 °C.

**Figure 8:**
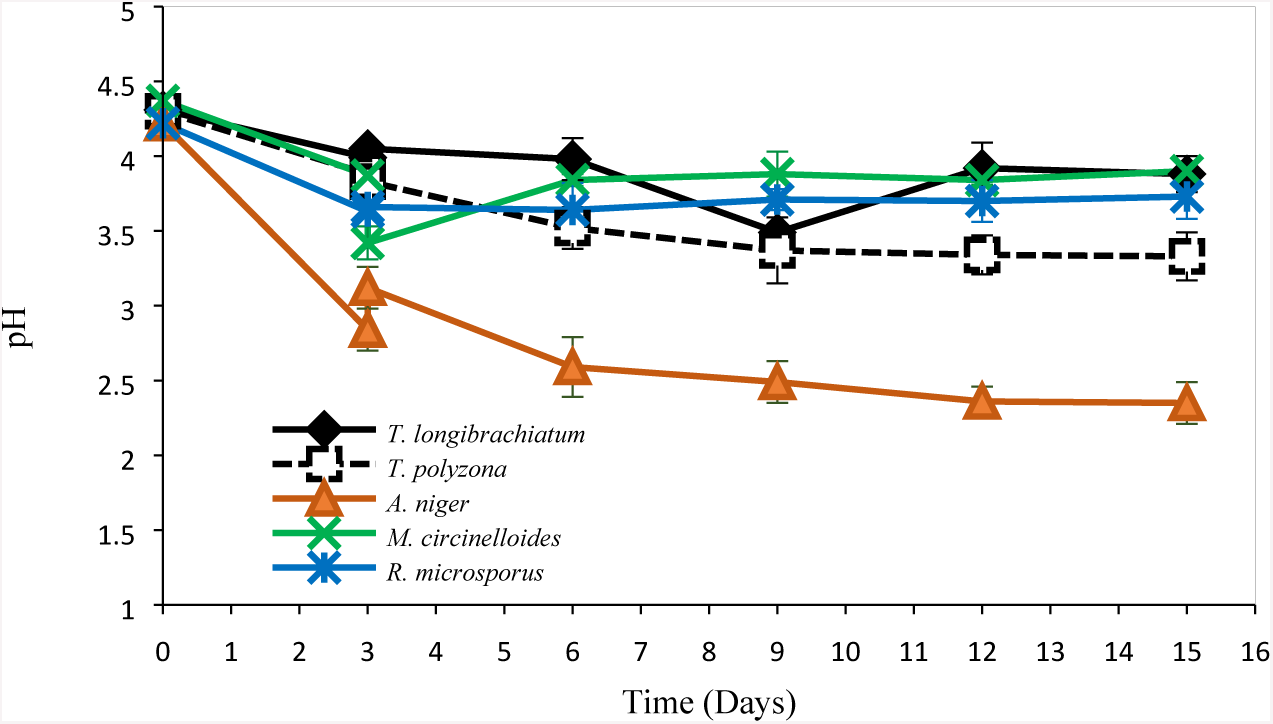
Evolution of pH in LN-m at 30±1.5°C.

From the starting pH, day 0 to the 5 day‘s values in LN-m during fungal mycelium formation, all the observed pH values were acid. This might be due to the fact that fungi produce a large range of fine enzymes and biochemicals including acid in suspension medium. Therefore, the pH of fungal suspension media were found to be acid regardless of the starting pH, this unlike bacteria that evolved in alkaline media. Figure 8 showed the evolution of the pH in fungal mycelium for 15 d from the initial pH of 4.3. The values represent the average of 3 flasks and error, the standard deviations. There was no significant different noticed on the final pH, of the 5 d of mycelium incubation, except between *T. polyzona* and *A. niger* (F=11.302 and *p* = 0.0099< 0.05: Figure 8).

The pH of fungal mycelium was found to be all acid. From the starting pH of 4.3, the pH values of the liquid media were decreasing and were found in the range of 2.4 to 4. Fungi are growing in acid media according to their secretion (Figure 8) unlike, bacteria which prefer an alkaline medium. The lowest pH values of 2.35±0.12 were found in the batch containing *A. niger.* Thus, significant difference *p* < 0.05 was noticed from *A. niger* flask compare to others fungal flasks, while no major difference was perceived within the rests. This low pH values unfavourable for bacterial growth and some pathogen proliferation, might be attributed to the high concentration of acid production from this fungal strain.

### Fungal growth in batch flasks

The following Figure 9 gave the set up of stationary batch flasks without daylight (Figure 9A), stationary and aerated batch flasks at the starting day (Figure 9B) and the aerated batch flasks after 15 d (Figure 9C). The aerated batch flasks were supplied with humidified air through DDW flasks.

**Figure 9:**
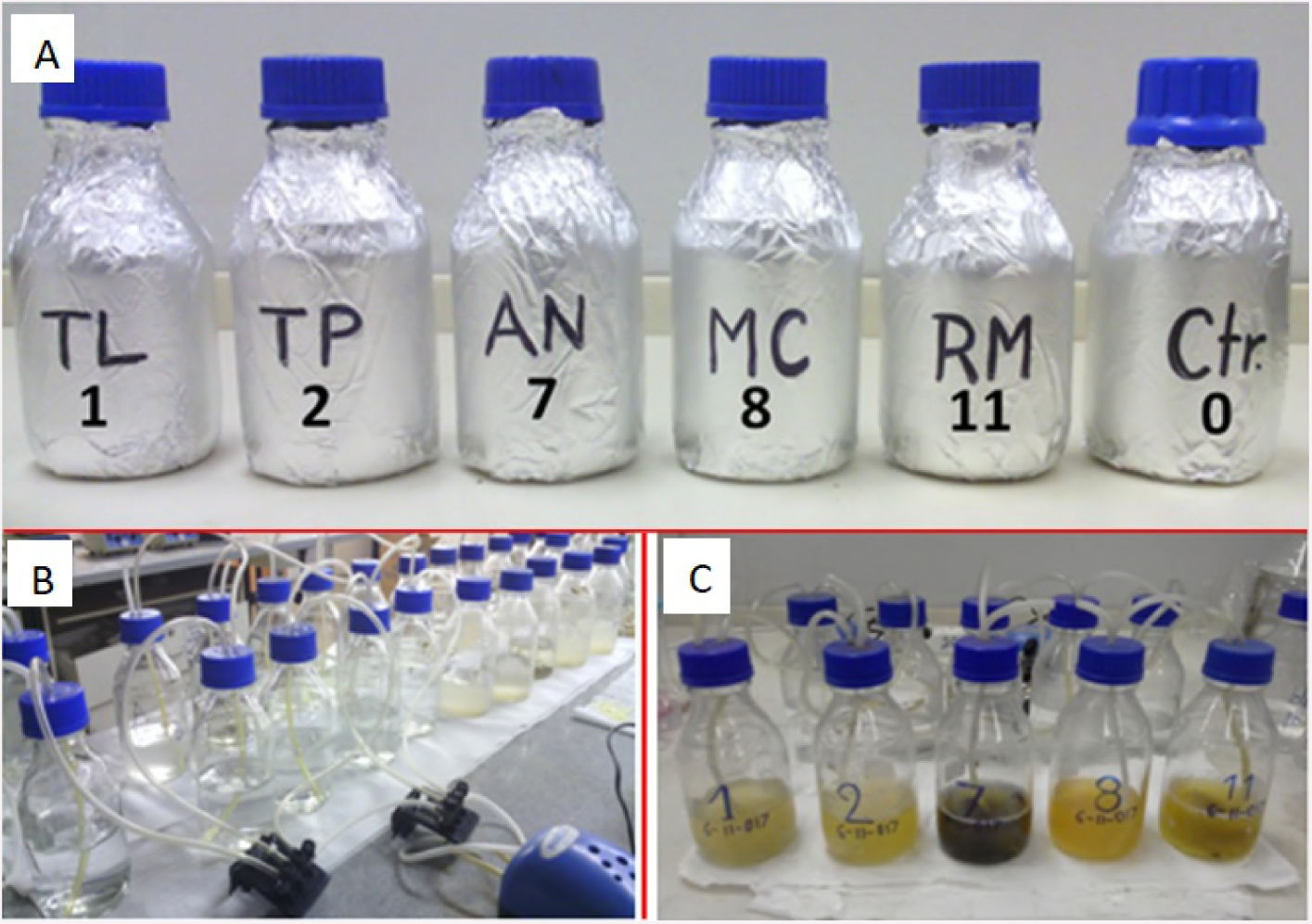
Stationary (StBF) and aerated (ABF) batch flasks set up (with TL or 1: *T. longibrachiatum,* TP or 2: *T. polyzona,* AN or 7: *A. niger,* MC or 8: *M. circinelloides,* RM or 11: *R. microsporus* and Ctr or 0: the control batch flask)

#### Effect of air supply in batch flasks

The fungal growth study was performed in 500 mL flask containing 200 mL of medium as batch flasks for 15 d at room temperature. Therefore, fungal growth experiments were carried out in StBF and in ABF as shown in Figures 10 and 11 below. The type of batch flask used (Figures 10 and 9) was found to affect significantly the fungal growth (*p* < 0.01). Although the fungal biomass appeared to be lower in ABF compare to the one in StBF, the air supply seemed typically to favour continual fungal mycelial growth (Figure 11). Most of fungi still growing after 9 and 12 d in ABF, while they start decreasing (dying) from the 6 and 9 d in StBF except *A. niger* which start at 12 d. However, the *T. longibrachiatum* biomass was not significantly affected by the growth conditions: its biomass appears to be in the same range (*p* > 0.5), while *T. polyzona* fungal cells showed good growth in ABF than in StBF. Its cells start dying and fungal biomass decreasing from day 6 in StBF, whereas it still growing in aerated batches. Even though *A. niger* gave the best growth in batch flasks as noticed in LN-m above regardless of working conditions (Figures 10 and 11). However, *A. niger* was growing from day 3 (41.23±3.12 mg/200mL) to the day 9 (52.55±8.15 mg/200mL) and day 15 (75.25±15.12 mg/200mL) in stationary batch flask and from day 3 (30.99±2.9 mg/200mL) to the day 9 (41.01±3.25 mg/200mL) and day 15 (55.65±4.05 mg/200mL) in aerated batch. *A. niger* and *R. microsporus* were found both to be affected by the batch conditions.

**Figure 10:**
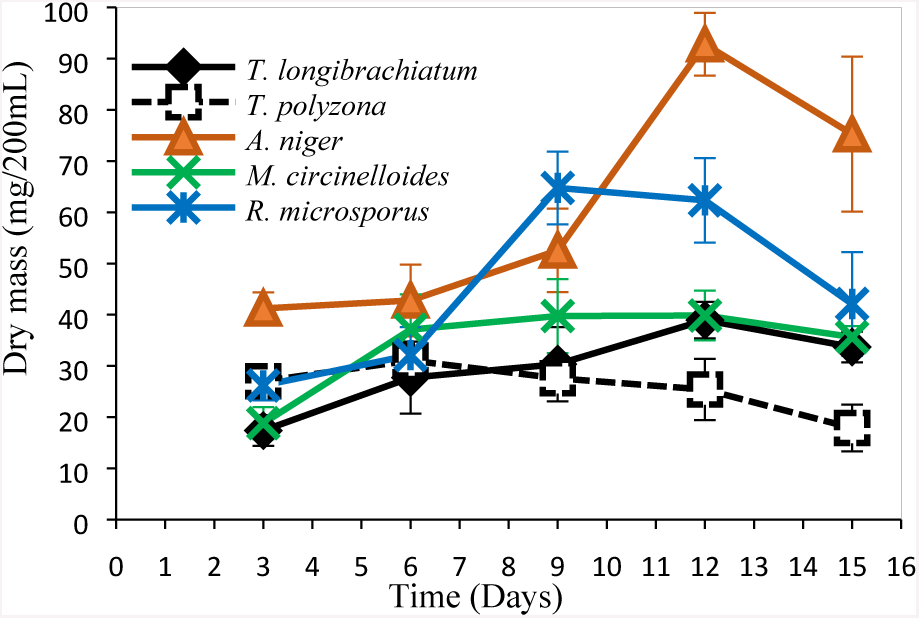
Growth study in StBF (10g/L glucose)

**Figure 11:**
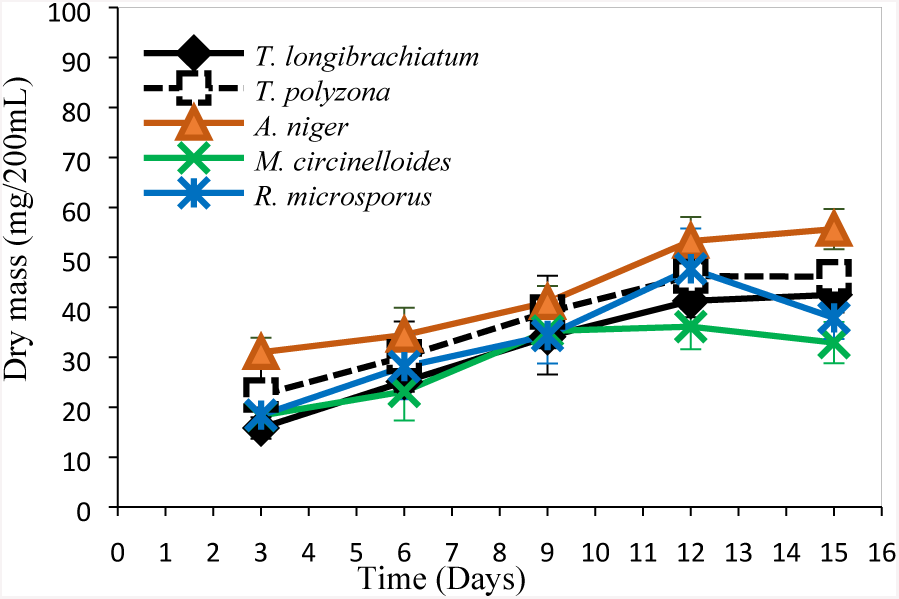
Growth study in ABF (10g/L glucose.

#### Effect of nutrients and daylight

The effects of nutrients (glucose as carbon source and ammoniun tartrate as nitrogen source) as well as daylight were conducted in StBF as expressed in the following Figures 12, 13, 14 and 15 below. The glucose concentration has affected significantly the fungal biomass growth (*p* < 0.001) (). Particularly *A. niger* showed in Figure 12 when compare to Figure 10 above (with dry mass recorded for example at day 3: 22.89±1.99mg/200mL against 41.23±3.12 mg/200mL and at day 12: 30.98±3.48 versus 92.81±6.12 mg/200mL). However, a slight increase was recorded in fungal dry mass due to the supplementation of AT in batch solution (Figures 10 compared to 14 and Figures 11 compared to 15 respectively in StBF and ABF). The AT supply seemed not to affect significantly the fungal growth (*p* = 0.321 > 0.05 in the StBF and *p* = 0.213 > 0.05 in ABF). Nevertheless, it might be noticed that, the batch flask without glucose nor AT should not be considered as not containing any nutrient. Indeed, the fungal inoculum from LN-m contains already macro and micronutrients for which the impact in fungal growth. Despite the fact that nutrients have been diluted in batch media, it‘s enough to sustain fungi for a certain period of time. *A. niger* looked affectively affected by glucose nutrient supply contrary to *R. microsporus*. By comparing the Figure 11 to Figure 8, the daylight appeared to affect a bit the fungal growth in stationary batch flask (*p*=0.201>0.05). Nevertheless, lower fungal biomass was recorded in absence of light (Figure 10 compare to Figure 13), absence of glucose (Figure 10 compare to Figure 12) and AT (Figure 10 compare to Figure 14 and Figure 11 to Figure 15).

**Figure 12:**
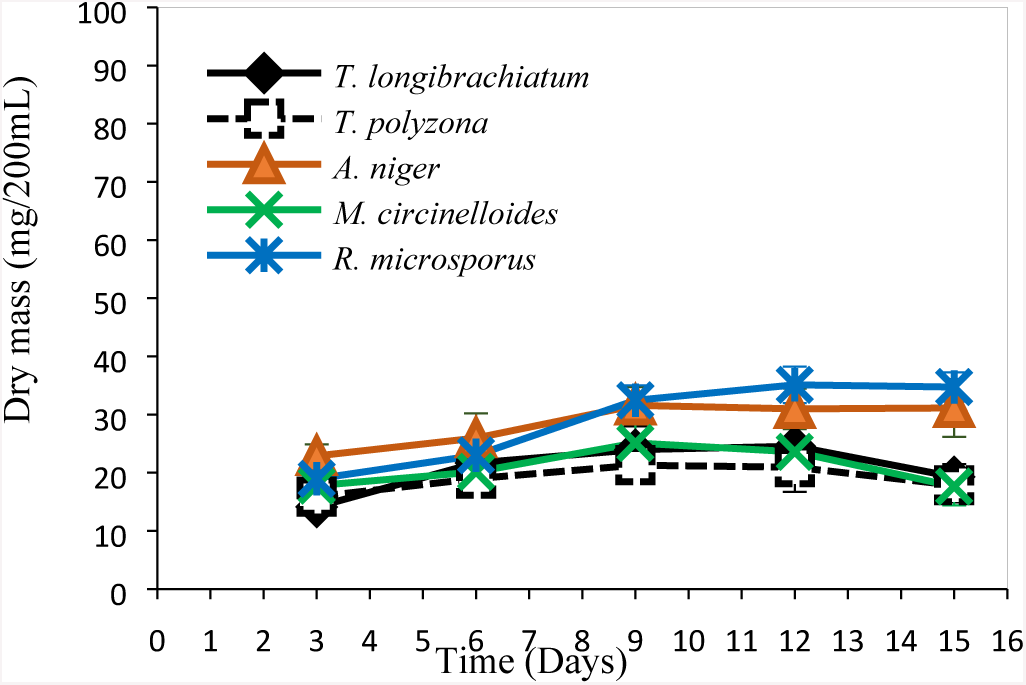
Growth study in StBF (0g/L glucose)

**Figure 13:**
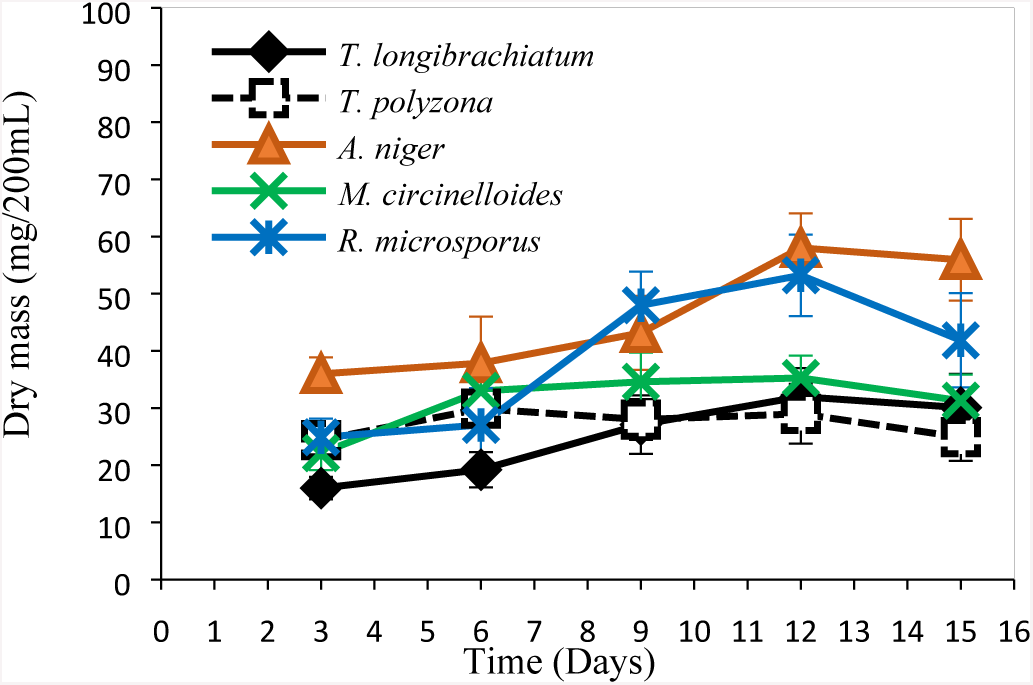
Growth study in StBF (10g/L glucose) without light.

**Figure 14:**
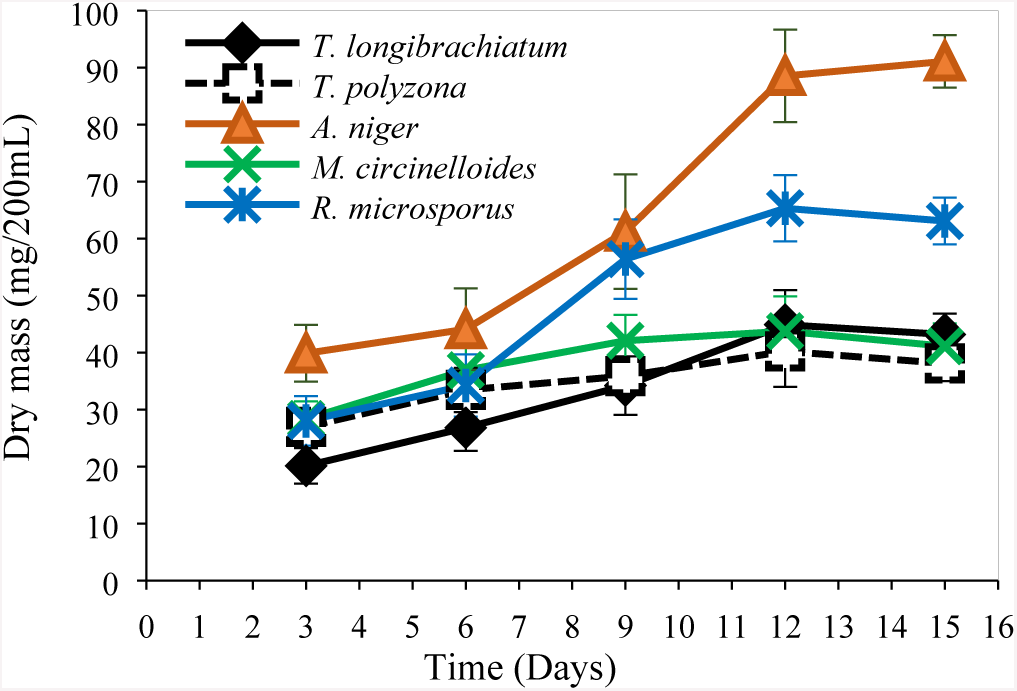
Growth study in StBF (10g/L glucose + 0.2g/L AT)

**Figure 15:**
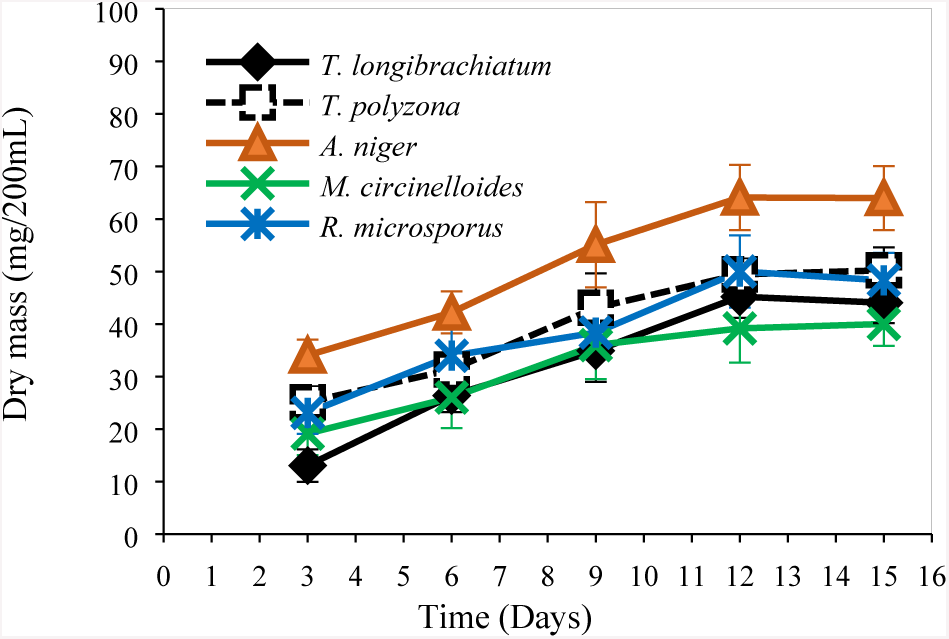
Growth study in ABF (10g/L glucose + 0.2g/L AT)

#### Effect of sterile conditions

The non-sterile conditions in StBF (Figures 16 against 10) did not affect significantly the fungal biomass (*p* = 0.83 >0.05). For example, at day 3 and day 12, the recorded dry mass were respectively found to be: for *A. niger* 36.44±4.67 mg/200mL against 41.23±3.12mg/200mL and 85.21±11.09mg/200mL against 92.8±6.12 mg/200mL, for *M. circinelloides* 20.05±4.09 mg/200mL against 18.83±3.13 mg/200mL and 33.09±10.01 mg/200mL versus 39.85±4.85 mg/200mL, for *T. polyzona* 26.19±2.87 mg/200mL against 27.03±1.99 mg/200mL and 27.63±4.62 mg/200mL versus 25.42±5.99 mg/200mL, for *T. longibrachiatum* 14.12±3.05 mg/200mL against 17.4±2.99 mg/200 mL and 35.59±8.19 mg/200mL versus 38.95±3.55 mg/200mL and then for *R. microsporus* 29.09±3.81 mg/200mL against 26.21±2.9 mg/200mL and 59.63±7.11 mg/200mL versus 62.33±8.25 mg/200mL. Nevertheless, lower fungal biomass was recorded in non-sterile batch flasks (Figure 10 compare to Figure 16).

**Figure 16:**
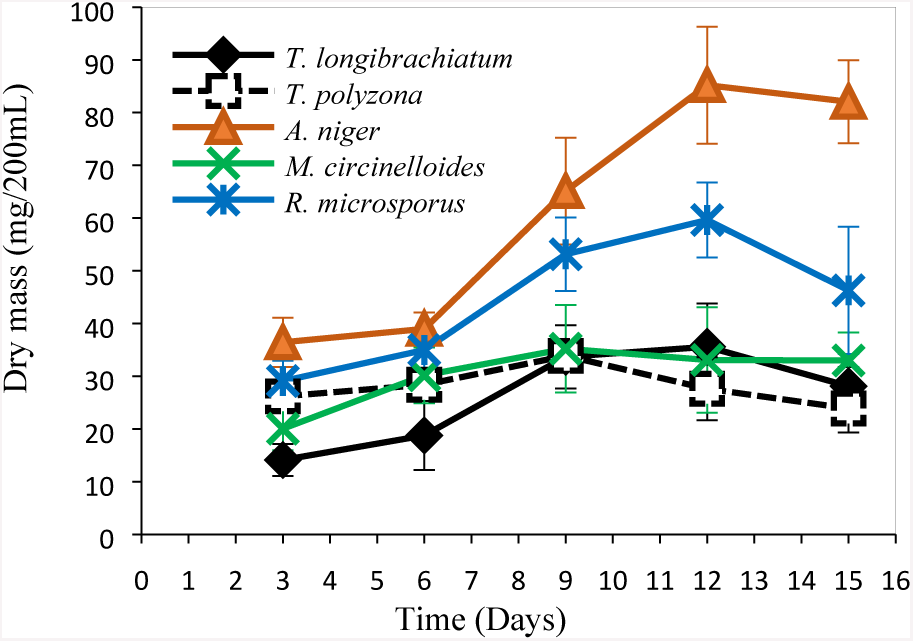
Growth study in non-sterile StBF (10g/L glucose)

### Diclofenac sodium removal from batch flasks

To ascertain the ISAIFS degradation efficiency of DCF contaminated synthetic wastewater, the experiments were performed in ABF and StBF. The DCF removal efficiency was assessed using the UV-Vis spectrophotometric method based on the reaction between DCF and concentrated nitric acid 55% (2:1 v/v).

#### Effect of nutrient, light and non-sterile conditions in DCF removal

The decreasing of DCF concentrations in batch flasks at different working conditions was thereby conducted in respect of its removal effectiveness by indigenous fungal strains achieved ABF and in StBF as showed by following Figures 17 to 22. The gradual decrease of DCF concentrations was recorded in general, except in the ABF control (Figure 17). The fungal removal efficiency was found to be better in ABF for *M. circinelloides, T. longibrachiatum* and for *R. microsporus* while compare to the StBF which gave good removal for *A. niger* and *T. polyzona.* The highest DCF removal was achieved by *R. microsporus* (95% after 7 d with pH 4.2±1) in ABF (Figure 17), whereas the lowest was by *T. longibrachiatum* (17.8%, pH 4.1) in StBF (Figure 18). However, from the first day, more than 50% of DCF removal was achieved by *R. microsporus* and *A. niger* in ABF, when less than 30% was recorded in StBF. The DCF removal efficiency reached in fungal consortium batch in StBF was not such a good, compare to individual performance.

**Figure 17:**
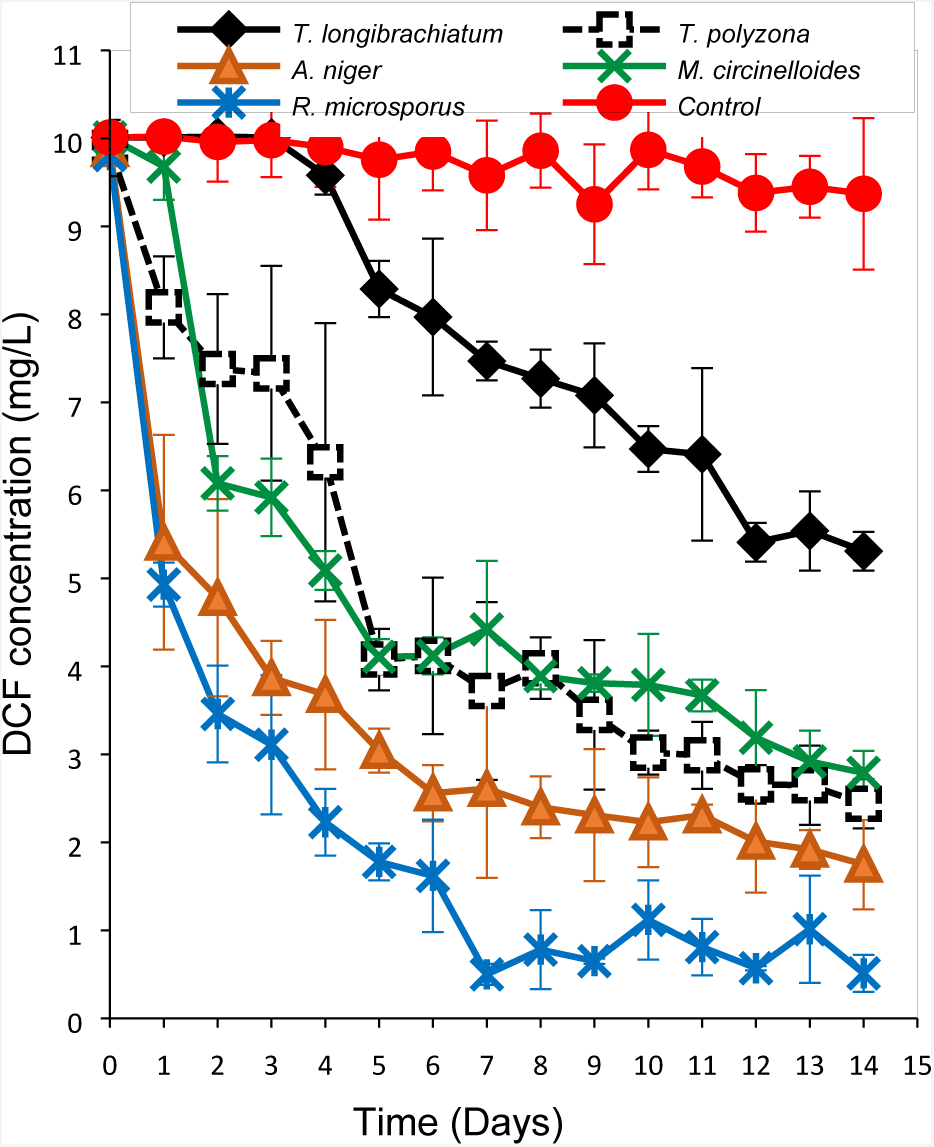
DCF removal in ABF (10mg/L of glucose)

**Figure 18:**
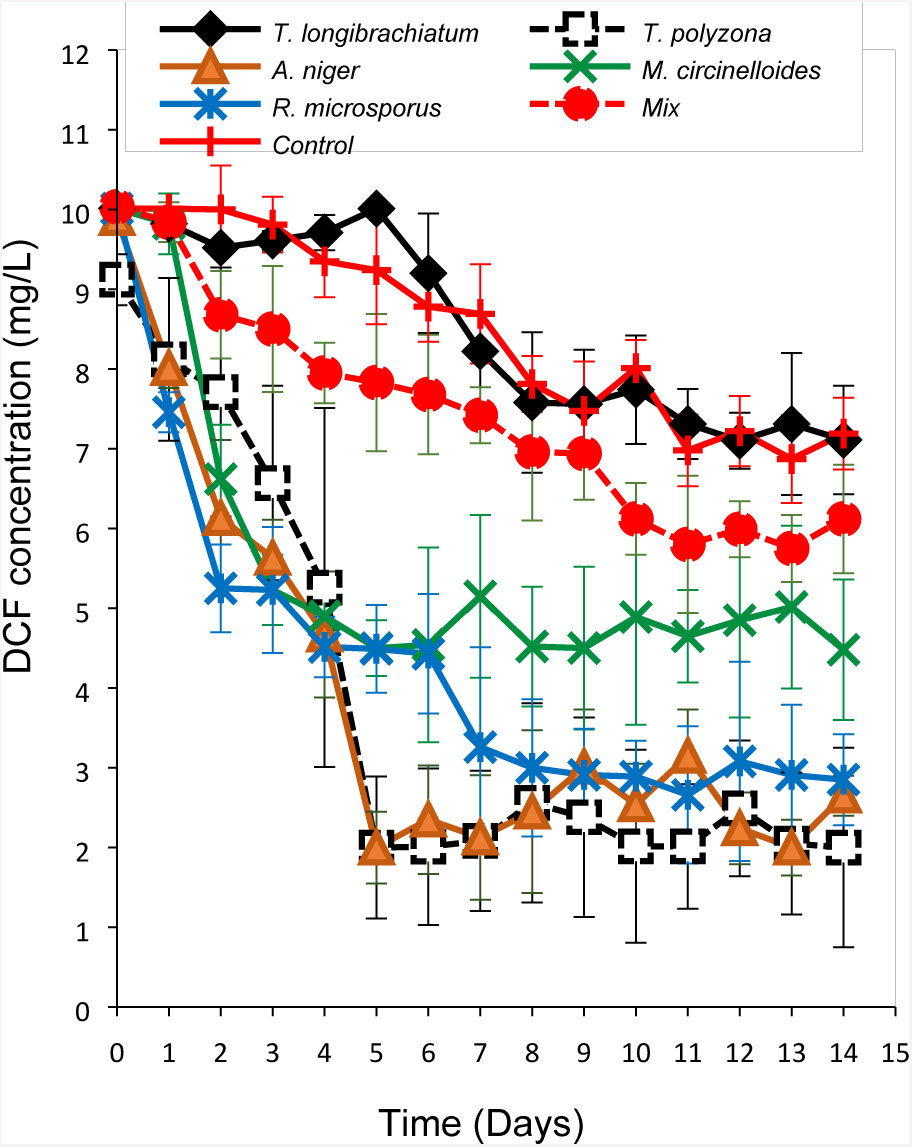
DCF removal in sterile StBF (10mg/L of glucose) with *Mix*= fungal consortium.

The removal efficiencies of the five ISAIFS in ABF (after 5 d) were as follows: 17.12% for *T. longibrachiatum*, 59.22% for *T. polyzona*, 69.58% for *A. niger*, 58.89% for *M. circinelloides* and 82.21% for *R. microsporus.* A lowest DCF removal was noticed in the ABF containing *T. longibrachiatum*. After 5 d in the sterile StBF, the Figure 18 showed: 0% for *T. longibrachiatum*, 80% for *T. polyzona*, 80% for *A. niger*, 55% for *M. circinelloides* and 55.1% for *R. microsporus*. At this Lab scale, no fungal synergic significant positive effect of mixed strains was noted in the DCF removal efficiency (Figure 18).

Despite the fact that after 2 d almost 50% and 40% of DCF removal were already reached respectively by *R. microsporus* and *A. niger*, the best removal was obained after 5 d in the StBF using *T. polyzona* and *A. niger* followed by *M. circinelloides* and by *R. microsporus.* However, none DCF removal should be noticed with *T. longibrachiatum* in StBF.

A slight degradation of DCF was noticed in each ABF control. This might be due to the photodegradation effect of DCF in aqueous solution reported by previous studies. However, this was accentuated in StBF (±25%) than in ABF. Compared each others, the ABF was found to give a good DCF removal in general, even to the point of attaining 17.12% of DCF removal (after 5 d) for *T. longibrachiatum,* and then followed by the StBF. Hence, the DCF removal was lowest *T. longibrachiatum* regardless of the batch flask. Furthermore, *T. polyzona* gives a good removal in StBF than in aerated flasks ABF. However, *R. microsporus* and *A. niger* seemed to give better DCF removal regardless of the flask used.

However, in order to evaluate the fungal capability to withstand harsh conditions (without nutrients), the StBF were performed with 0 mg/L of glucose (Figures 19 and 21). The non-sterile batch StBF (Figures 20 and 21) were performed against the sterile batch StBF (Figures 18, 19 and 22). The daylight effect was also conducted in sterile StBF (Figure 22).

**Figure 19:**
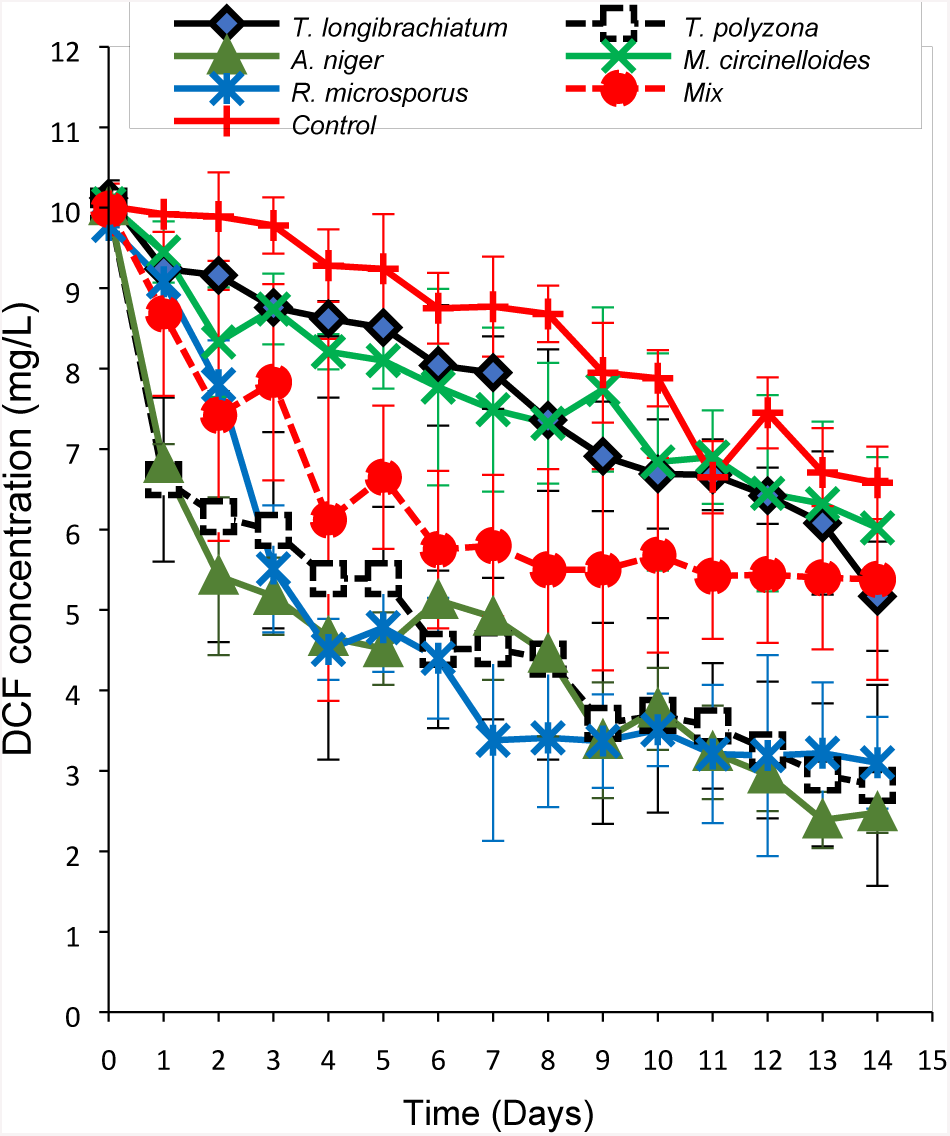
DCF removal in sterile StBF (0 mg/L of glucose)

**Figure 20:**
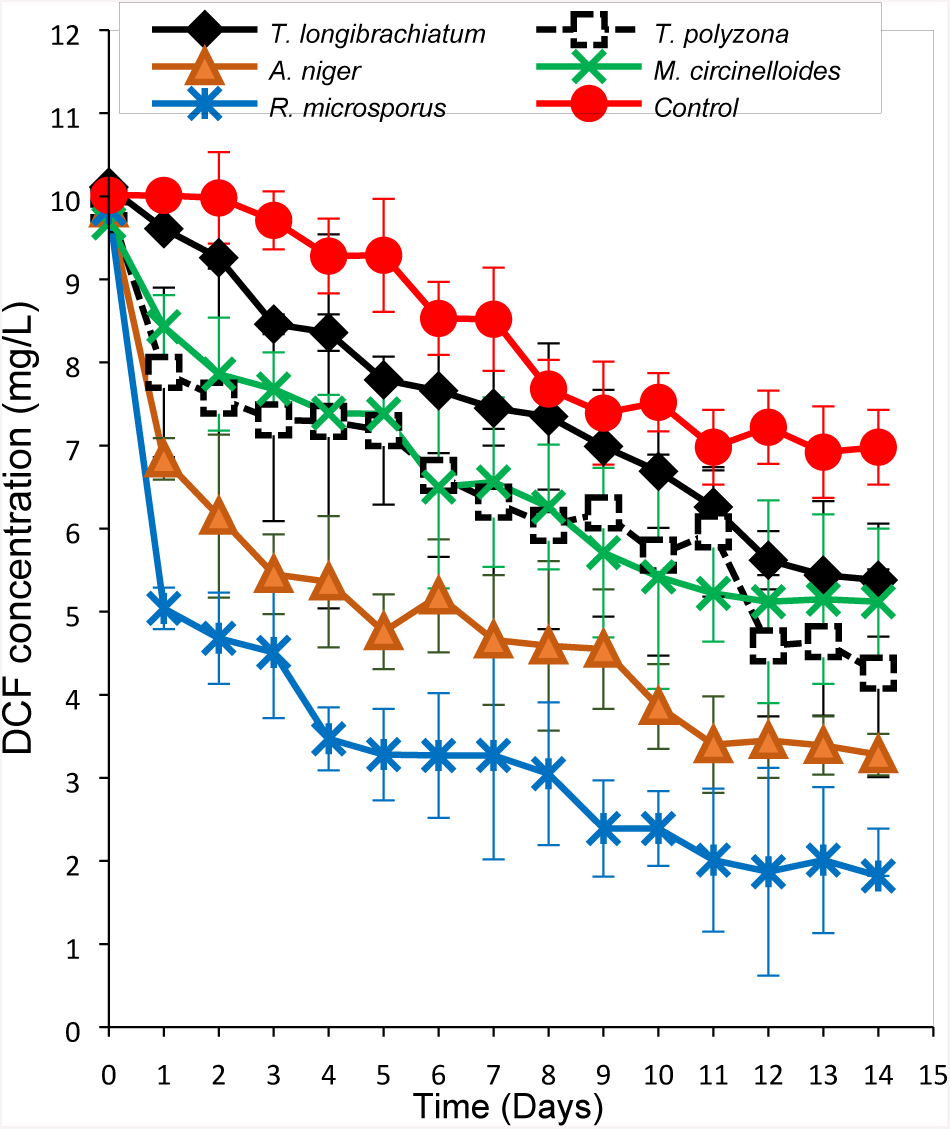
DCF removal in non-sterile StBF (10 mg/L of glucose)

**Figure 21:**
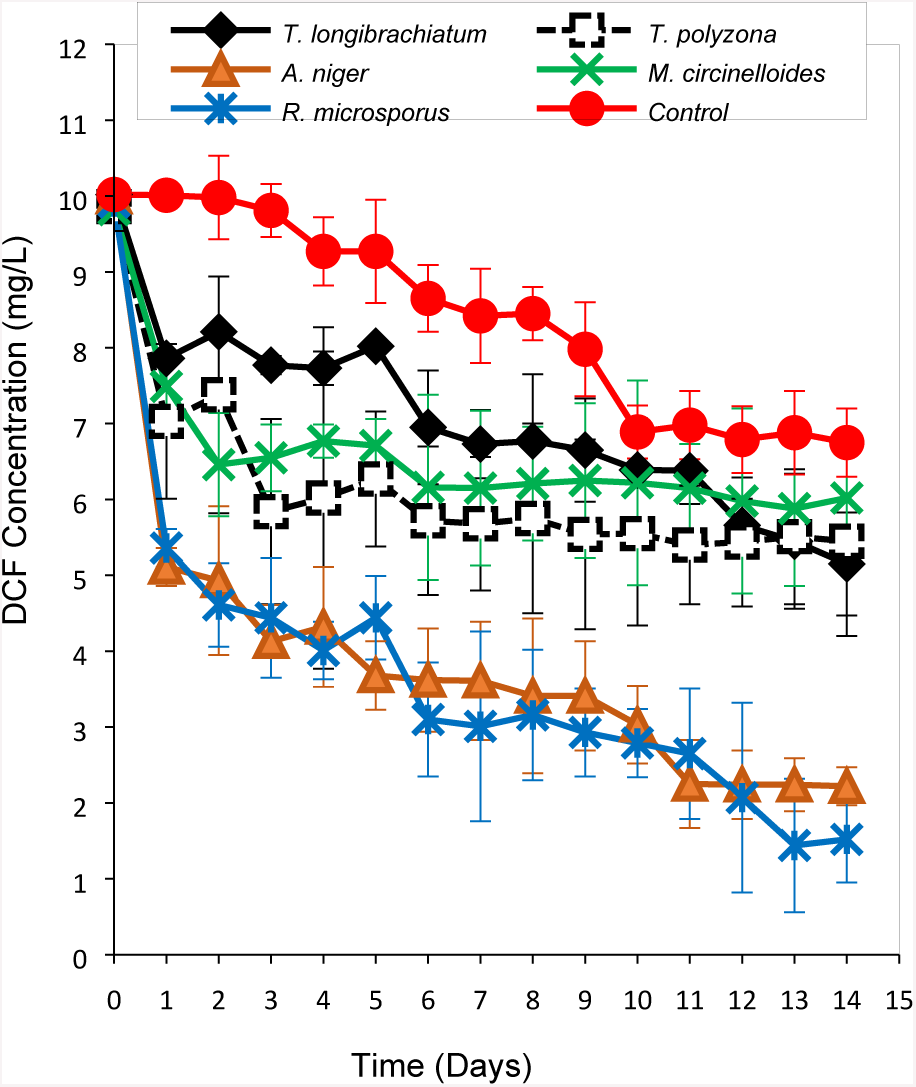
DCF removal in non-sterile StBF (0 mg/L of glucose)

**Figure 22:**
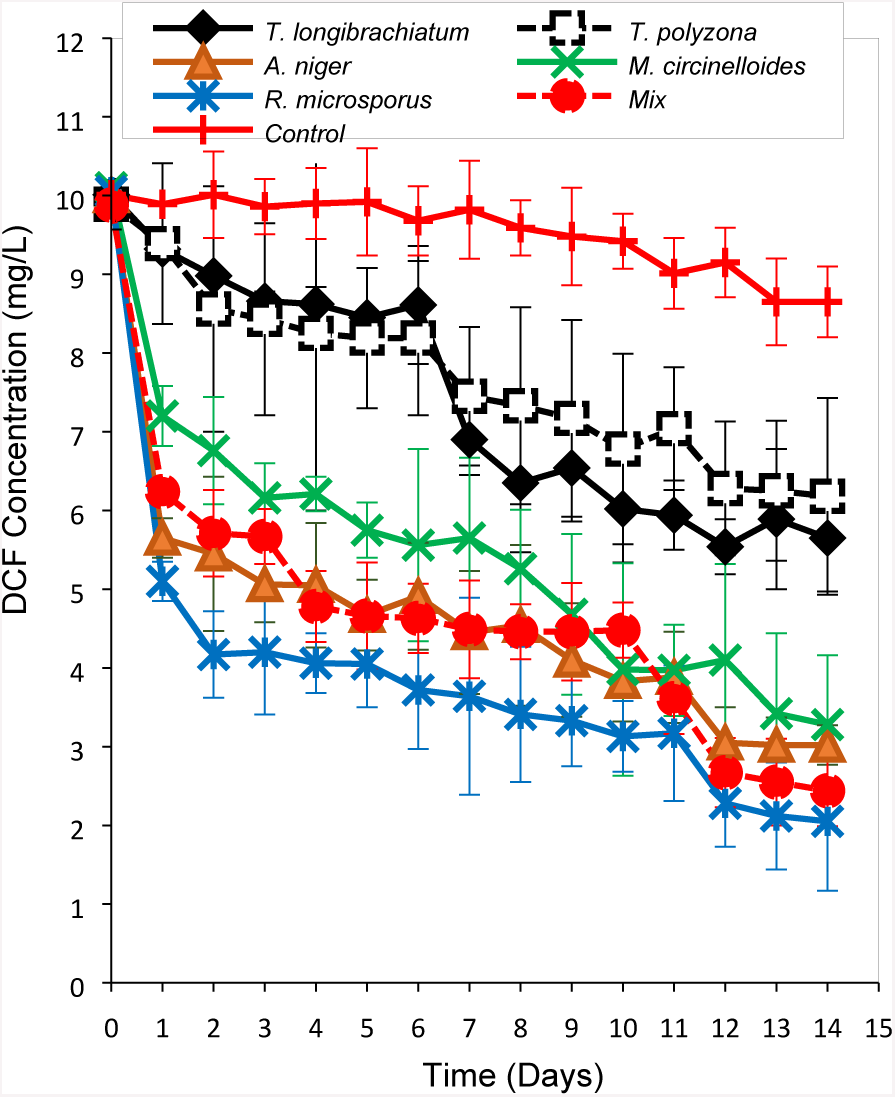
DCF removal without daylight and sterile StBF (10mg/L of glucose)

The lack of glucose in StBF seemed to reduce the DCF removal efficiency (Figure 20 compare to Figure 19) and affected the fungal growth (Figure 10 compare to 12). After 5 d, ± 80% of DCF removal was reached by both *T. polyzona* and *A. niger* in sterile StBF supplied with glucose (Figure 18), whereas only ±46.1% and ±54.8% were respectively achieved in non glucose supplied batches (Figure 19). However, the fungal consortium was found to give better DCF removal in absence of glucose (±21.7% in Figure 18 and ±33.5% in Figure 19). The removal efficiency of *T. polyzona, M. circinelloides* and *A. niger* appeared to be significantly affected by the absence of nutrient (*p <* 0.05), while *R. microsporus* seemed not to be considerably affected (from ±55.1% to 52.2% of DCF removal).

In addition, *R. microsporus* performed better in non sterile batches than in sterile conditions (±67.2% and ±55.1% after 5 d: Figures 18 and 20). Furthermore, the DCF removal ability of *T. polyzona* and *A. niger* were negatively affected by non-sterile conditions [from ±80% of DCF removal efficiency for both (Figure 18) to ±28.2% and 52.4% respectively (Figure 20)]. Consequently, the none glucose supply in non-sterile StBF did not significantly change the DCF removal ability of *R. microsporus* (*p >* 0.05), whereas the *A. niger* was positively affected (Figure 21 compare to Figure 19). Its DCF removal was found to be good in StBF without glucose (Figure 21) than in the rests of StBF (Figures 19, 20 and 22).

The negative effect of daylight in DCF removal was noticed in general in StBF (Figure 18 compare to Figure 22). The removal efficiency of *T. polyzona* and *A. niger* were negatively affected in absence of daylight, while it appeared to enhance the DCF elimination efficacy of fungal consortium. Furthermore, by comparing Figure 18 to Figure 22, the DCF photodegradation noticed in Figure 18, disappeared in the absence of light (Figure 22).

#### Effect of fungal biomass concentration in DCF removal

The DCF removal efficiency seemed to be dependent on the mycelium inoculum concentration as showed in Figure 23. The increase of fungal biomass from 5% to 30% of mycelium concentration in StBF affected the DCF removal. Hence, the fungal removal efficiencies were increased by 7.9% for *T. longibrachiatum*, by 20.4% for *T. polyzona*, by 30% for *A. niger*, by 19.7% for *M. circinelloides* and by 2.72% for *R. microsporus.* Although the continual increase in fungal mycelium inoculum concentration seemed to enhance DCF removal efficiency of *T. polyzona* and of *T. longibrachiatum,* there is no real effect observed for all fungal strains beyond 30% in the experimental conditions (Figure 23). Regardless of the mycelium concentration, *R. microsporus* gave a high DCF removal, followed by *A. niger* and lowest removal recorded for *T. longibrachiatim.* However, the fungal mycelium concentration increase seemed to favor the basidiomycetes WRF *T. polyzona*‘s removal efficiency.

**Figure 23:**
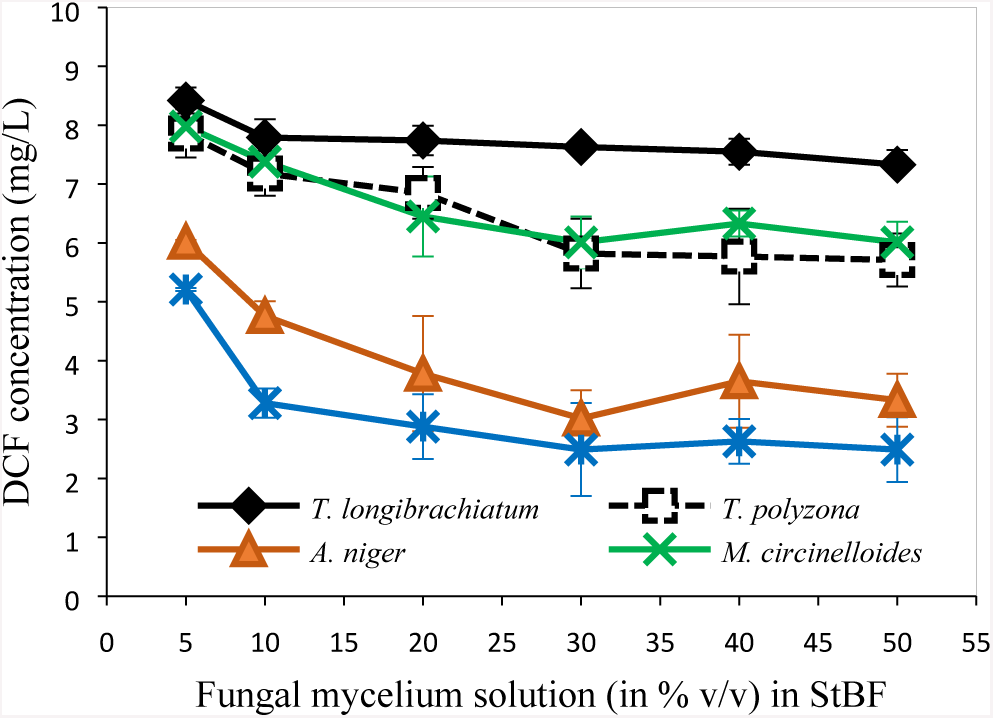
Fungal mycelium concentration effect in DCF removal from StBF.

#### Possible fungal DCF degradation pathway

Previous studies have reported that, the DCF degradation process went through an unknown mechanism which ended up by the releasing of CO_2_. The following Figure 24 showed the unidentified mechanism pathway suggested by Fabregat (2014).

**Figure 24:**
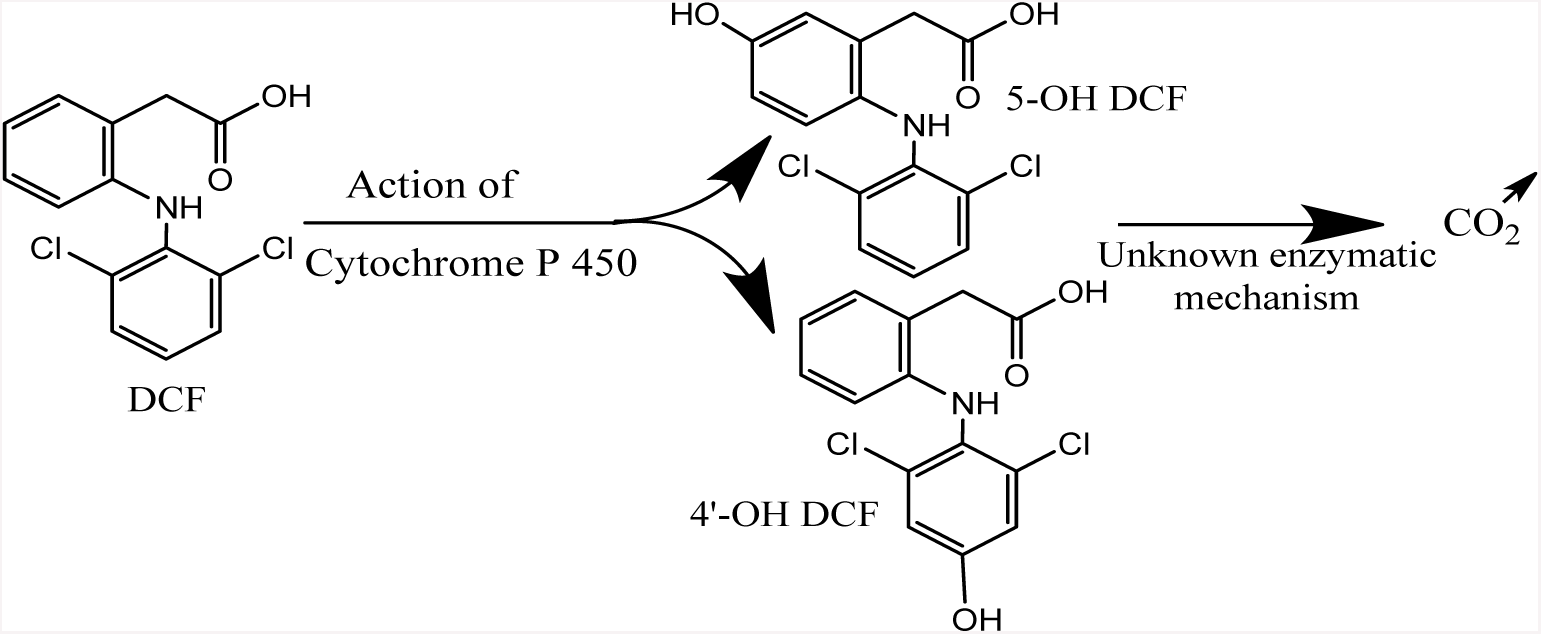
Degradation pathway of DCF by *T. versicolor*, where intermediates: 4‘-OH DCF = 4‘-hydroxydiclofenac and 5-OH DCF = 5-hydroxidiclofenac.

### Statistic analysis

The initial pH of the fungal mycelium cultures in LN-m, seemed to affect their growth. From the Figure 6 above, the following R-boxplots (Figure 25a and 25b) illustrated the pH effect within and between isolate fungi. Regardless of the initial pH of LN-m, *A. niger* gave a better growth compare to others (*p <* 0.001) (Figure 25a), followed by *R. microsporus* and *M. circinelloides*. The lowest growth was recorded with *T. longibrachiatum.* However, no significant pH effect in fungal growth within fungal strains was noticed (*p* = 0.123 > 0.05, Figure 25b).

**Figure 25:**
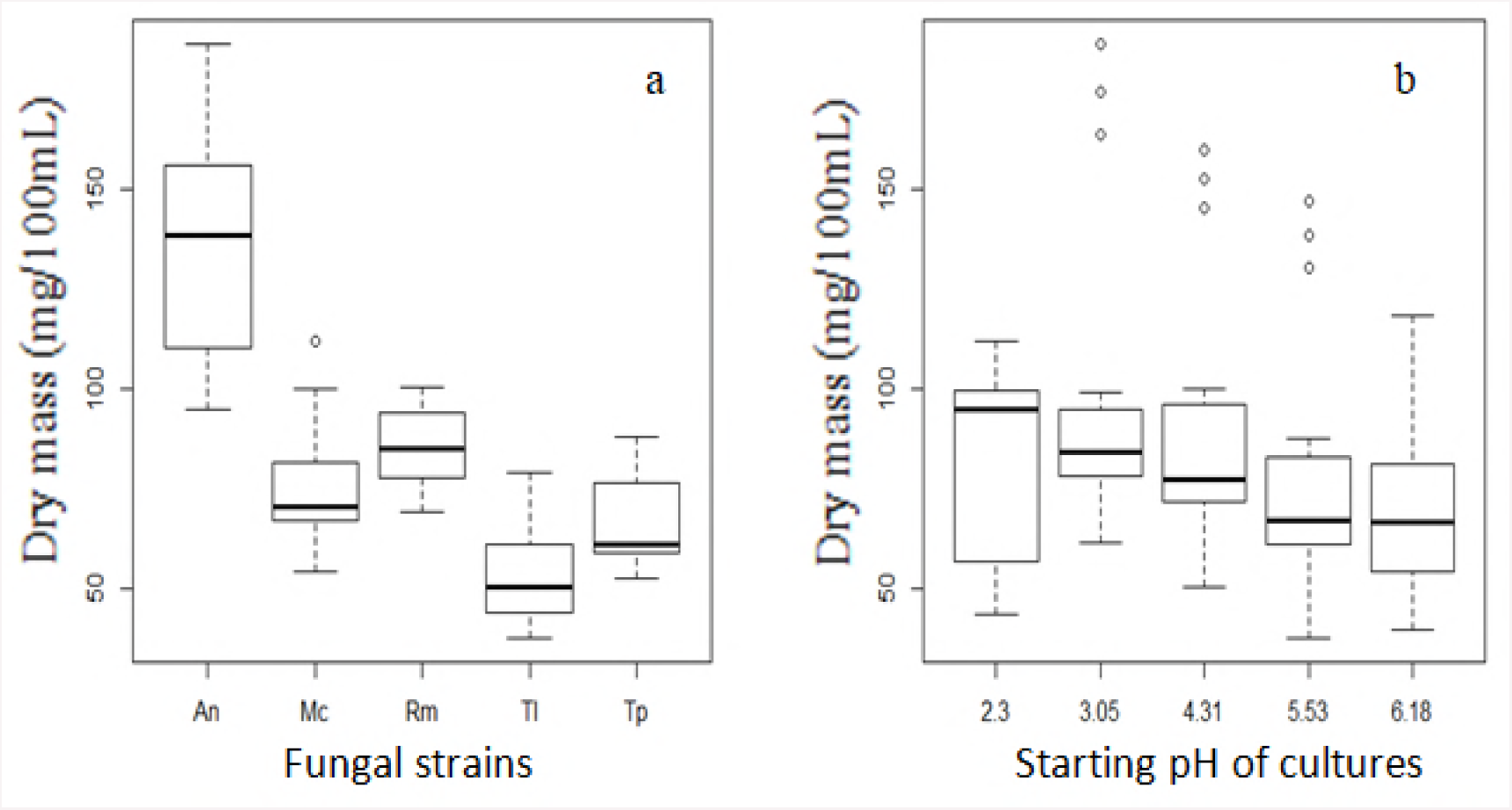
a. Boxplot of pH effect between fungi and b. Boxplot of pH effect within fungi. Where, An: *A. niger*, Co: Control, Mc: *M. circinelloides*, Rm: *R. microsporus*,Tl:*T. longibrachiatum*, Tp: *T. polyzona* and Mix: mixture batch containing all 5 fungi.

In addition, from Figure 5 above, the following R-boxplots (Figures 26a and 26b) showed the effect of temperature in fungal growth. The strain *A. niger* showed a significant thermotolerance to the increase of the temperature (*p <* 0.001) when compare to others (Figure 26a). The fungal growth in suspension liquid media was temperature dependent (Figure 26b). A negative correlation of the increasing of the temperature on fungal growth was noticed with the coefficients: −0.994, −0924, −0.992, −0.975 and −0.968 respectively for the five ISAIFS: *T. longibrachiatum, T. polyzona, A. niger, M. circinelloides* and *R. microsporus* (Figure 26b)

**Figure 26:**
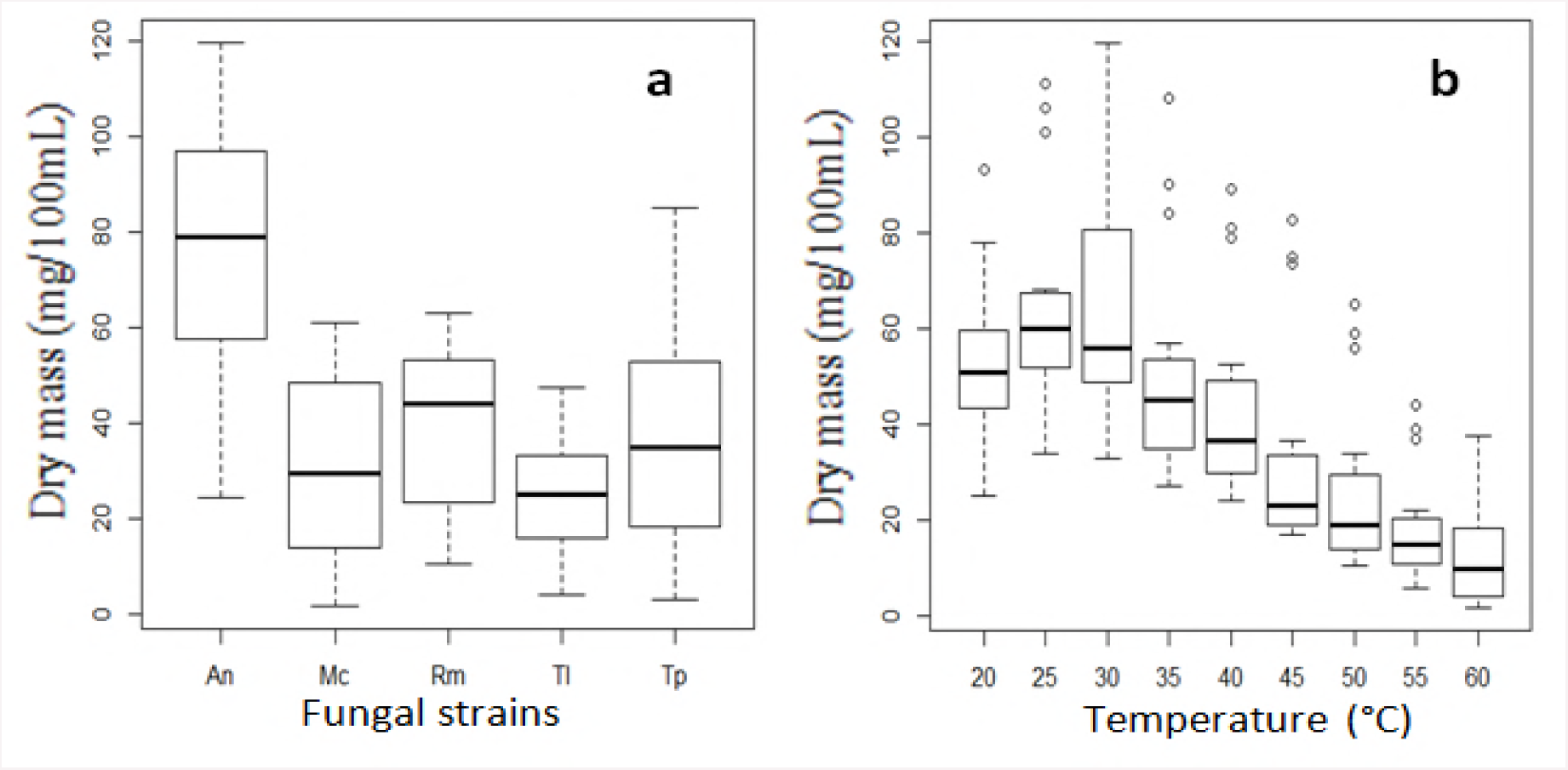
a. Boxplot of temperature effect between fungi and b. Boxplot of the effect within fungi.

The fungal DCF removal efficiencies reported in the following Figure 27 (R-boxplot of DCF removal in %) were calculated from Figures 17 to 22 above. It can be seen from Figure 27 that, comparing to the control and regardless of working conditions, ascomycetes (*A. niger* and *T. longibrachiatum*), azygomycetes (*M. circinelloides* and *R. microsporus*) and basidiomycetes (*T. polyzona*) indigenous fungi had a certain DCF removal capabilities at different levels. This was reflecting the DCF oncentration decreasing recorded from Figures 17 to 22. The air supply (ABF: Figures 17 and 27a: continuous air supply compared to StBF: Figures 18 and 27b: limited air supply) affected negatively the removal efficiency of *A*. *niger* and *T. polyzona* (*p <* 0.05), while it favoured *R. microsporus* and *T. longibrachiatum* removal. *M. circinelloides* was not affected. The lack of glucose nutrient (Figures 18 and 27b compare to Figure 19 and 27c) affected significantly the fungal removal efficiency (*p* < 0.001) regardless of species in sterile conditions. However, while performed in non-sterile conditions (Figures 20 and 27d compare to Figures 21 and 27e), the absence of glucose seemed in general not to affect significantly the removal of fungi (*p* > 0.05).

**Figure 27:**
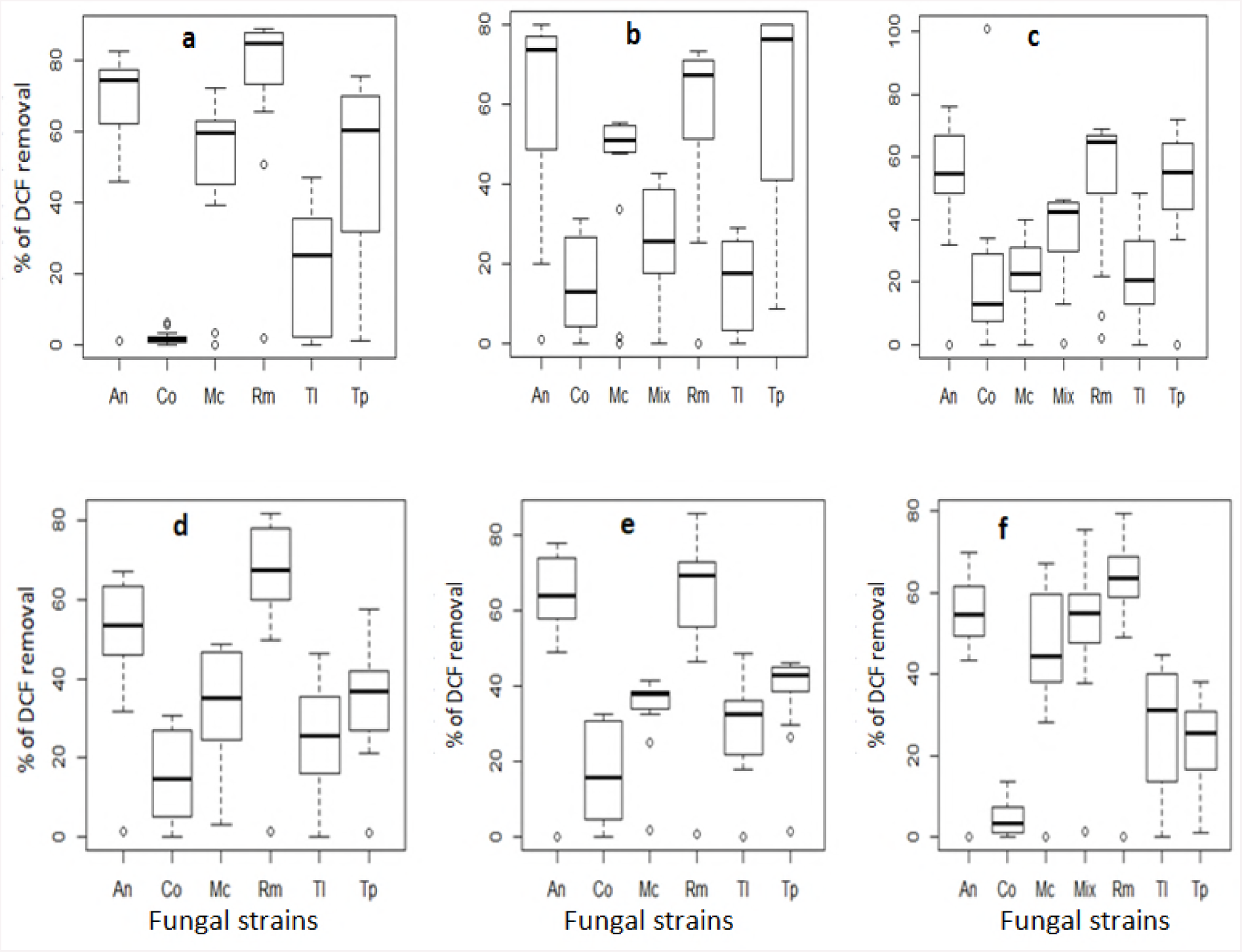
Boxplot of DCF removal, a: ABF (from Figure 17), b: sterile StBF (from Figure 18), c: sterile StBF (from Figure 19), d: non-sterile StBF (from Figure 20), e: non-sterile StBF (from Figure 21), f: sterile StBF (from Figure 22).

The sterile operating conditions (Figures 18 and 27b compare to Figures 20 and 27d) favored significantly the fungal DCF removal efficiency in glucose supply batches (*p <* 0.05). However, in absence of glucose (Figures 19 and 27c compare to Figures 21 and 27e) the non-sterile conditions affected positively the removal (*p* < 0.05), especially for *A. niger, M. circinelloides, R. microsporus* and *T. polyzona*. The daylight effect (Figures 18 and 27b compare to Figures 22 and 27e) seemed to favour the DCF removal efficiency (*p* < 0.05) regardless of fungal species.

The following Figure 28 showed the effect of the quantity (%) of fungal mycelium inoculated in StBF in the DCF removal efficiency. The removal of DCF in StBF is significantly improved with the increase of fungal mycelium from 5 to 30% (*p* = 0.0013 < 0.01). After 30% of fungal mycelium, its increase seemed not the influence the DCF removal in StBF in our experimental conditions (*p* > 0.05) (Figure 28a). Even though the fungal biomass was increased, the lowest removal was recorded for *T. longibrachiatum,* whereas *R. microsporus* continued to give the best DCF removal, followed by *A. niger.* This increase seemed to enhance a bit the *T. polyzona* removal efficiency (Figure 28b).

**Figure 28:**
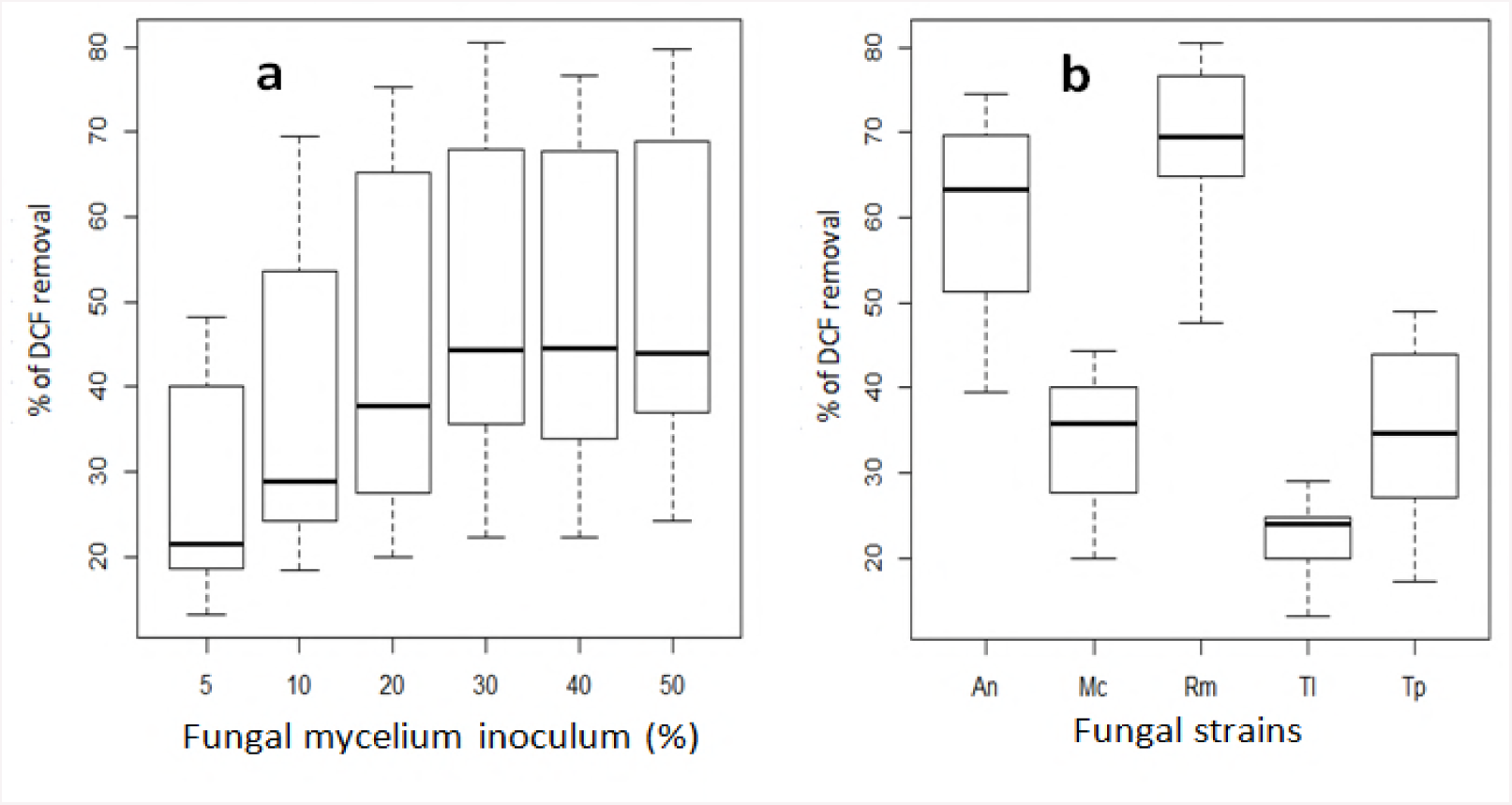
Boxplot of DCF removal (%), a: % of fungal mycelium inoculated in StBF, b: effect of inoculum % in DCF removal efficiency of fungal strains (from Figure 23)

## Discussion

Currently, pharmaceutical compounds are among the emerging organic pollutants in wastewater and aquatic environment that has raised significant concern (Yang et al., 2017). In general, among the most abundant pharmaceuticals found in wastewater and surface water, they are the non-steroidal, analgesic and anti-inflammatory drugs such as acetaminophen, diclofenac, ibuprofen and naproxen (Rivera-Jaimes *et al.,* 2018). Although some of them showed a certain removal throughout WWTPs, their concentrations downstream were found to be only slightly lower than upstream. Hence justifies their presence into the river. In addition, the concentrations of diclofenac, ibuprofen, naproxen and sulfamethoxazole present in the river might cause a high toxicity risk for the aquatic ecosystem (Rivera-Jaimes *et al.,* 2018). Considering the continuous introduction of pharmaceuticals into the environment with possible transformation into multiple and most often unknown byproducts, they may have the harmful effect into wildlife and human health. Consequently, the unwanted biological effects may occur on the exposed organisms, due to the fact that pharmaceuticals byproducts have been found sometimes to be more toxic and persistent than the parent compounds (Noguera-Oviedo and Aga, 2016; Rivera-Jaimes *et al.,* 2018). Because the pharmaceutical‘s ecotoxicological impacts in the environment is not yet subject to any regulation (Rivera-Jaimes *et al.,* 2018).

The present study investigates the isolate fungal growth optimization in suspension liquid media and evaluates their DCF removal efficiency from synthetic wastewater. Therefore, different working conditions were explored in order to optimize the indigenous fungal degradation of DCF contaminated synthetic wastewater.

In general, an increase on fungal growth was observed by considering the huge changes noticed in fungal morphology and size growth from the third day. That would be due to the spores swelling process based on the continuous diffusion of water and nutrients over the spore surface as described by Grünberger *et al*. (2017) to its germination and hyphal extension for branch initiation of mycelium formation. In addition, there is a normal and almost random distribution of fungal growth noticed in the following 9 d of incubation in 100mL flasks suspension liquid media, which might be due to the continuously growing spores and mycelium proliferation. At the meantime, some fungal cells are dying and releasing of its cytoplasm into the media as reported by Grunberger *et al.* (2017).

The nitrogen requirement of ISAIFS showed that AT was the best nitrogen source, when compare to peptone as reported by Dutta and Das (2017). Brian *et al*. have also been shown since 1947 that, while glucose is used as carbon source, fungus (*Murothecium verrucaria*) grew readily with AT as nitrogen source. However, the carbon/nitrogen ratio in liquid media seemed to promote significant fungal growth (*p* < 0.001). The C/N ratio in liquid medium (LN-m, 50/1) is different and high from the one used by Jonathan and Fasidi (2001), (2/3 for *Psathyerella atroumbonada,* Pegter). Hence, for the majority of fungi, ammonium salts look to be a suitable source of nitrogen, especially AT as reported by Dutta and Das (2017). The value of *p* < 0.001 shows the significant effect of media in fungal growth, with better growth in the medium with high C/N ratio (LN-m). Although PL-m has previously recorded a higher fungal biomass by using peptone as nitrogen source with lactose as carbon source, low fungal growth was recorded by using peptone with glucose anhydrous. No significant different was noticed in isolate fungal growth (*p >* 0.05), when ammonium sulfate (AS) was used as nitrogen source with glucose anhydrous as carbon source.

Besides, among nutrients, dead cells constitute potential nutrient for living cells (Fu and Viraraghavan, 2000, 2002). This could give a prediction of a plateau or stability of fungal growth and decay curve, which would justify the fungal biomass recovery and reuse in batch flasks for EDCs removal.

The nutrients supply (glucose and AT) and light seem to affect significantly the fungal growth in batch flasks. Additionally, according to Webster and Weber (2007), light is an essential parameter for fungal sporulation, stimulation of conidial development as well as the growth of sporangiophores, even though it doesn‘t affect zygospore formation. Light rouses aerial fungal growth from hyphal bodies. Therefore, switching to larger cell aggregates resulting in greater fungal object tightly connected pixels due to the lack of fresh medium supply during 14 or 15 d of experiment. Indeed, according to Grünberger *et al.* (2017) and Espinosa-Ortiz *et al.* (2016), the pellet morphology on fungal growth could be supported by the continuous infused fresh medium through the system in order to establish constant cultivation conditions. According to Rodarte-Morales *et al*. (2012b), the fungal pellet are formed in submerged batch media, especially in aerated batch flask, while in stationary batch flask, fungal filament shaped mycelium grow and invade the batch media volume.

Furthermore, the temperature of the study shows that, the range of 27°C to 31.5°C is suitable for fungal growth regardless of strain. Fungal biomass recorded was less at 37±1°C. At this temperature, the duration (day) is also a considerable parameter effecting fungal biomass. Although other fungal strains started dying from around the day 12, *A. niger* grew quickly and also still growing even beyond 15 d. This confirms the remarkable resistance of *Aspergillus* species (ubiquitous fungal saprophytes able to face diverse ecological niches worldwide) even in harsh conditions as reported by Garcia-Rubio *et al*. (2017). This quality might be used for repeated fungal batch flasks as shown by Lu *et al*. (2016). The isolate *A. niger* strain is equally well endowed with these qualities. However, Zhang and Geißen, (2010) reported that, the temperature was been found to be an important parameter which affect even the EDC degradation efficiency of *P. chrysosporium* strain BKM-1767 (DSM 6909). In addition, the fungal enzyme such as lignin peroxidase, responsible of pollutants biodegradation was found to be temperature dependence (Tien and Kirk, 1988; Zhang and Geißen, 2010). Hence, 30±1.5°C seems to be optimum temperature for fungal growth. This value was proposed to suitable for fungal enzymes production (Kumar *et al.*, 2016; Teerapatsakul *et al.,* 2017). Therefore, by using *T. polyzona* as thermotolerante fungi, Teerapatsakul *et al.* (2017) have found that when temperature was increased from 30°C to 37°C and 42°C, only laccase and manganese peroxidase were produced during the biodegradation process of polycyclic aromatic hydrocarbons. This shows the effects of the temperature in fungal metabolites production.

The isolate *R. microsporus* did not show any thermotolerance in its growth until 45°C, while *R. microsporus* isolate by Dolatabadi *et al.* (2014) has been considered as good candidate for the thermostable amylase enzyme production. Dolatabadi *et al*. (2014) have found in their study that, the suitable temperature for *R. microsporus* was between 36°C and 45°C, with an optimum at 45°C. According to van Leeuwen *et al*. (2012), fungal strains are considered to be mesophile fungi because of its thriving at moderate temperature of about 20±1°C to 40°C. Therefore, the isolate South African indigenous fungi can be considered as mesophile fungi.

The range of pH of 3.1 to 5.5 is found to be favourable for fungal growth. This range included the optimum of pH 4.3 recommended for enzymes activity (Tien and Kirk, 1988; Zhang and Geißen, 2010; van Leeuwen *et al.*, 2012). Despite the pH variation, *A.niger* has given, a highest fungal growth showing its strange resistance to harsh conditions, followed in order by *R. microsporus, M. circinelloides, T. polyzona* and *T. logibrachiatum.* However, the VA supply at the third day to stimulate enzymes production, was found to affect a little bite the pH. The optimum pH for most fungal strain media is found to be in the acid range of pH < 5.0, where enzymatic activity seems maximal as well as fungal metabolism such as availability of cell permeability and biochemical production such as citric acid (Koduri and Tien, 1994; Zhang and Geißen, 2010; van Leeuwen *et al.*, 2012; Ranawat and Rawat, 2017). In addition, the structure and composition of cytoplasmic membrane in cells can be also denatured that defines the substrate consumption rate of fungi (van Leeuwen *et al.*, 2012; Puig, 2012). The decreasing of pH noticed in fungal media will not allow bacterial proliferation in fungal liquid culture (Asokan *et al*., 2016). Furthermore, the fungal DCF removal efficiency in general is related to its enzyme activities (Zhang and Geißen, 2010). DCF has been completely degraded in the pH range of 3.0 - 4.5 by Zhang and Geißen (2010) using crude lignin peroxidase from *Phanerochaete chrisosporium*. In the present study, the pH of fungal submerged liquid culture (fungal mycelial generation in LN-m and fungal growth in batch flasks) was found to decrease from the starting pH 4.3. This might due to the substrate consumption especially when AT is used as nitrogen source as well as metabolite production (Haapala *et al.,* 1994; Asokan *et al*., 2016).

Even though fungi are growing faster in StBF (with air limited) than in ARF (with continuous air supply), their biomass are decreasing earlier from day 6 (for *T. polyzona*) and day 9 (*R. microsporus*) in StBF, whereas it happens later from day 12 in ARF. At the meantime, *R. microsporus,* and *T. polyzona* was found to still growing over 15 d. This means that fungi were rapidly growing in StBF as well as they were speedily dying. The ABF appears to be a suitable fungal batch flask.

Referring to previous investigators, the similar of isolate fungi were used for different purposes. For example, *A. niger* was used in the simultaneous fermentation and saccharification of ethanol in co-culture with *Saccharomycea cerevisiae* (Izmirlioglu and Demirci, 2017), in the production of citric acid (Wang *et al.,* 2017) and in the degradation of 85% of Azo dyes Reactive Red 120 (Ameen and Alshehrei, 2017) as well as 98.5% of Congo Red in immobilization mycelial pellets batches (Lu *et al.,* 2017). However, up to date, no study has been conducted in the removal of DCF. Therefore, in this study the DCF removal efficiency was reached by *A. niger* batches at 73.89±10.14% in ABF and 78.75±7.8% in StBF after 7 d.

Although the strain *M. circinelloides* isolate by Giraud *et al.,* (2001) was not found to be efficient while used in the elimination of polycyclic aromatic hydrocarbons especially anthracen and fluoranthene (Giraud *et al.,* 2001), the South African indigenous isolate strain *M. circinelloides* gives after 5 d respectively 58.9±2% and 55±3.5% of DCF removal efficiency in ABF and StBF. Furthermore, the similar strain used by Mirbagheri *et al.* (2016) has found to be useful to enhance the alpha-linolenic acid production in the fermentation of industrial oil wastes.

On the other hand, the isolate strain *R. microsporus,* achieves a good DCF removal about 95±1.2% in ABF and 73.4±8.6% in StBF respectively within 7 and 11 d. Nevertheless, the strains *R. microsporus* and *A. niger* isolated by Bordjiba *et al.* (2001) were not found to be sensitive in the degradation of herbicides contaminants (triazines: metribuzin and metamitron, phenylureas: linuron and metobromuron). The isolate fungus *T. polyzona* classified among WRF by Gauthier *et al.* (2017) gives a DCF removal of 69.79±2.5% after 10 d in ABF and 80±8.9% within 5 d in StBF. The similar WRF strain has shown its biodegradation efficiency in textile reactive dyes decolorization: anthracene (Remazol Brilliant Blue R) and phthalocyanine (Synozol Turquoise Blue) at 97% and 80% respectively (Cerron *et al.,* 2015) as well as in bisphenol A biodegradation (Chairin *et al.,* 2013).

Even if certain studies have shown the efficacy of fungal consortium, for example the WRF (*Phenerochaete chrysosporium, Trametes versicolor* and *Fomes fomentarius*) performed to enhance the properties of the final compost product from municipal waste after 37 d by Voberskova *et al.* (2017), the isolate fungal consortium (Mix batch) conducted in sterile StBF seems to give low DCF removal rate compare to individual fungal batches. The consortium fungal DCF removal of 42±4.5% is recorded after 11 d, better than the one from *T. longibrachiatum* alone.

However, considering the initial DCF concentration of few ng/L to 15μg/L found in wastewater (Li *et al.*, 2014; Verlicchi *et al.*, 2015), after up to 95% of its removal efficiency in our experimental study, the pollutant issue might be sorted out. The DCF concentration will decrease to less than 7.5μg/L and 0.75μg/L respectively after 1 and 5 d. In addition, knowing that isolated fungi growth in acid media, they can be used for the treatment of pharmaceutical industry wastewater, which has an acid pH as reported by Gadipelly *et al.* (2014).

Preceding investigations have reported that DCF has been removed after only two hours at 77% by performing an oxygenated batch with *Phanerochaete chrysosporium* (Rodarte-Morales *et al.*, 2012b) and for total removal with almost complete elimination of its acute lethal toxicity towards the freshwater after 6 d using *Phanerochaete sordida* YK-624 enzyme (Hata *et al*., 2010), whereas a total removal has been completed by lignin peroxidase (Zhang and Geißen, 2010) and by laccase (Lloret *et al.*, 2010) in 1 h. Nevertheless, the DCF removal effectiveness reached in this study was better than the one conducted in a nitrifying-denitrifying plant (less than 10%) after 1 d (Suárez *et al.*, 2005), in membrane bioreactor (10 - 45%) (Reif *et al.*, 2008) and in the coagulation-flocculation and flotation process (20 to 70%) (Carballa *et al*., 2005).

In addition, several parameters were of non-negligible contribution in our experimental conditions for the purpose of DCF removal efficiency. First of all, the air supply plays an important role in fungal activity. According to Wells and Uota (1970), the air supply seemed typically favour fungal mycelial growth in liquid media. The living fungal cells needed molecular oxygen for their respiration. The availability of oxygen may affect the consumption of carbon and nitrogen in the media. However, it has been reported that ATP and ADP concentrations are significant in the control of glucose utilization and electron transport systems (Chance and Hollunger, 1961; Barker *et al*., 1964), knowing that the stimulation of respiration in low oxygen tension may affect a ATP decrease as well as change to an alternative electron transport pathway (Harrison and Pirt, 1967) or the uncoupling of oxidative phosphorylation (Racker, 1965).

The nutrient sources (especially carbon and nitrogen) are cognitive factors for fungal germination and growth as well as metabolites production (van Leeuwen *et al*., 2012; Dutta and Das, 2017). Hence, as stated by Puig (2012) glucose was identified as the most appropriate substrate carbon source for fungal growth and highest production of manganese peroxidase enzyme in batch fermenter experiment, when associate with AT as nitrogen source. The use of AT has been reported to decrease the pH of submerged liquid culture of fungi (Haapala *et al.,* 1994). The DCF removal efficiency is higher in glucose supply batch flasks than in absence of glucose (*p* < 0.05), especially for *A. niger, M. circinelloides, R. microsporus* and *T. polyzona*. Therefore, the removal efficiency hereby is slightly higher for *A. niger* and *T. polyzona* than the one observed for *R. microsporus,* which was good in absence of glucose. To the high fungal biomass recorded in glucose supply batches corresponded to a good DCF removal efficiency than its counterpart without glucose, where less fungal biomass was noticed. This might be due to relationship between fungal biomass and enzymatic activity. However, the continual increasing of fungal biomass in batch flask doesn‘t enhance continually the DCF removal effectiveness. This corroborate to the result found by Takey *et al.* (2014), where 2g/L of fungal biomass were giving best removal compare to 1g/L and 3g/L. The non-sterile conditions affect negatively the DCF removal efficiency of *A. niger* and *T. longibrachiatum*. The negative effect of non-sterile treatment on *A. niger* degradation efficiency was also reported by Pérez-Armendariz *et al.* (2010) during the removal of weathered hydrocarbons from soil of tropical Mexico. Furthermore, in non-sterile conditions, the lack of glucose supply doesn‘t affect the good DCF removal efficiency recorded for *A. niger, R. microsporus* and *M. circinelloides*, while *T. longibrachiatum* and *T. polyzona* were good in sterile batches, while no significant difference is noticed for *M. circinelloides* (*p* > 0.05). However, it has been reported that, the increase in biomass concentration can decrease the biosorption ability and affect the removal efficiency (Takey *et al.,* 2014).

The daylight effect has an additional effect in the DCF degradation. However, the effect of light in DCF phodegradation has been reported by Hashim *et al.* (2016) and Zhang *et al.* (2017), where the photocatalytic reactor was performed for DCF removal from wastewater.

Different DCF biodegradation pathways have been proposed. According to Tien and Kirk (1988), the fungal mycelium were collected at the fifth day to inoculate the batch flasks, while they still growing owing high fungal enzyme production. Accordingly, Zhang and Geißen (2010) have found that the optimum lignin peroxidase production occurred at the fifth day of *P. chrisosporium* culture. Previous studies have also attributed the removal of DCF as well as some recalcitrant pollutants including carbamazepine, to the intracellular cytochrome P450 enzyme system, in the *Trametes versicolor* culture. This DCF biodegradation went through an unknown mechanism which ended up by CO_2_ released (Marco-Urrea *et al.*, 2009, 2010; Fabregat, 2014). The openings found in hyphal branches are adsorption site of DCF as reported by Fu and Viraraghavan (2001, 2003) when removing micropollutants using dead fungal cells.

It has also been reported the biosorption and biodegradation as two fungal mechanisms for the removal of micropollutants from wastewater (Yang *et al.,* 2013). According to Fu and Viraraghavan (2002), the fungal biomass is regularly negatively charged on its surface. The negative polarity of functional groups plays an important role as biosorption sites of heavy metals and dyes by living and dead fungi (Fu and Viraraghavan, 2002). However, DCF sodium is an acid salt compound which releases an anionic ion with negative polarity after its dissociation from the sodium positive ion (Garcia-Lor *et al.,* 2010). Therefore, there is a predictable and possible repulsion between DCF and anionic group on the surface of fungal biomass. The possible fungal biodegradation pathway remains the enzymatic biodegradation (ten Have and Teunissen, 2001; Yang *et al.*, 2013).

## Conclusions

The fungal growth study in suspension liquid media has shown to be better in liquid media with high C/N ratio (50/1), with the temperature range between 26.5 °C to 31.5°C and pH from 3 to 4.5 considered as optimum conditions favouring fungal growth. *A. niger* has shown to growth faster and be temperature resistant than other ISAIFS. Then good dry mass were gotten stationary batch flask for *A. niger* and *R. microsporus* while *T. longibrachiatum* and *T. polyzona* have given the good growth in aerated batch flask, when *M. circinelloides* seemed to grow regardless of batch flask. ISAIFS seemed to need O_2_, glucose and AT nutrients supply and daylight for better growth in liquid media. The effects of glucose supply as well as light have been reported to play a significant role in fungal growth as well s in DCF removal efficiency. The non-sterile condition seemed to affect negatively the removal efficiency of some indigenous fungi.

The best DCF removal was achieved by *R. microsporus* (95% after 7 d) in an aerated batch flask at pH 4.2±1 and temperature (28±1°C). Although the fungus species mostly applied for the removal of trace organic compounds are *T. polyzona* (WRF) and *A. niger* this research indicated that several other species could also be used. This is the case of *M. circinelloides* and *R. microsporus.*

Our straightforward fungal flask systems which are based on the use of ISAIFS, seemed to present a certain capacity for DCF degradation in Laboratory scale. This appeared to be better than other traditional methods applied in WWTPs. It clearly shows that DCF biodegradation and removal from water might be attributed to the fungal enzymatic activities. Either lignin peroxidase and manganese peroxidase, laccase or cytochrom P450, the synergic effect of all fungal enzyme or metabolites would be the basis of the DCF fungal elimination efficiency.

This means that, it‘s possible to develop in wastewater treatment plants systems for recalcitrant pollutants removal using these indigenous South African fungal strains in the fungal flas for an ecological treatment of wastewaters. This is the first time that South African indigenous fungi especially *A. niger, M. circinelloides, T. longibrachiatum, T. polyzona* and *R. microsporus* are used for the removal of DCF from water and have shown a removal efficiency for this xenobiotic pollutant known as one of the makers for wastewater contamination by pharmaceuticals.

## Declarations

## Consent to publish

Not applicable

## Availability of data and materials

The data generated in this review, supplementary tables and figures are found in the supplementary file.

## Authors’ contributions

## Ethics approval

This article does not contain any studies concerned with experiment on human or animals.

